# Migratory behavior is positively associated with genetic diversity in butterflies

**DOI:** 10.1101/2022.06.21.496610

**Authors:** Aurora García-Berro, Venkat Talla, Roger Vila, Hong Kar Wai, Daria Shipilina, Kok Gan Chan, Naomi E. Pierce, Niclas Backström, Gerard Talavera

## Abstract

Migration is typically associated with risk and uncertainty at the population level, but little is known about its cost and benefit trade-offs at the species level. Migratory insects often exhibit strong demographic fluctuations due to local bottlenecks and outbreaks. Here, we use genomic data to investigate levels of heterozygosity and long-term population size dynamics in migratory insects, as an alternative to classical local and short-term approaches such as regional field monitoring. We analyze whole-genome sequences from 97 Lepidoptera species and show that migratory species have significantly higher levels of genome-wide heterozygosity, a proxy for effective population size, than non-migratory species. Also, we contribute whole-genome data for one of the most emblematic insect migratory species, the painted lady butterfly (*Vanessa cardui*), sampled across its worldwide distribution range. This species exhibits one of the highest levels of heterozygosity described in Lepidoptera (2.95% ± 0.15). Coalescent modeling (PSMC) shows historical demographic stability in *V. cardui*, and high effective population size estimates of 2 to 20 million individuals 10,000 years ago. The study reveals that the high risks associated with migration and local environmental fluctuations do not seem to decrease overall genetic diversity and demographic stability in migratory Lepidoptera. We propose a “compensatory” demographic model for migratory r-strategist organisms in which local bottlenecks are counterbalanced by reproductive success elsewhere within their typically large distributional ranges. Our findings highlight that the delimitations of populations are substantially different for sedentary and migratory insects, and that, in the latter, local and even regional field monitoring results may not reflect whole population dynamics. Genomic diversity patterns may elucidate key aspects of an insect’s migratory nature and population dynamics at large spatiotemporal scales.

## Introduction

Animal movement plays an essential role in configuring species’ population dynamics. Among different types of movement observed in nature, migratory behavior results in seasonal long-distance movements at the whole population scale (Dingle, 1996; Dingle & Drake, 2007; Jonzén et al, 2011). Quantifying the costs and benefits of migration is a challenge and, consequently, the trade-offs of this strategy are not well understood. A key question that remains is: What are the long-term net effects of migration on a species’ population size and genetic diversity? On the one hand, migration can be beneficial since it allows for rapid exploitation of alternative habitats in response to environmental changes (Chapman et al., 2015). On the other hand, migration is also associated with substantial risks that can lead to large fluctuations in offspring number between successive generations. Costs are typically caused by increased predation, exhaustion, disorientation and unpredictable weather during the migration, or unsuitable habitat at the destination, all of which can lead to high mortality compared to sedentary species, and thus severe decreases in population size or extinctions for particular migratory clusters (Newton, 2008; Wilcove & Wikelski, 2008; Calvert et al., 2009; Chapman et al., 2015; Møller, 2008; Saunders et al., 2019).

Migratory animals can also experience long-term population bottlenecks at the species’ range level. Since migratory ranges usually cover wide geographic areas, species can suffer from both long-term effects of climate change and habitat transformations in any of these areas, such as food shortage at critical breeding or stopover sites and/or increased exposure to diseases (Cox, 2010). It is therefore crucial to distinguish between bottlenecks/fluctuations that occur at a local and/or regional level, which are normally short-term, affecting individual migratory clusters within larger populations, and bottlenecks that occur at the whole population or species’ range level, which are usually spread over many generations and caused by dramatic events at a large spatial scale (England et al., 2003). Population declines have been observed in a range of migratory birds and other vertebrates (Sanderson et al., 2006; Berger et al., 2008; Moller et al., 2008; Both et al., 2010; Wilson et al., 2018), some of which have resulted in extinction (Hung et al., 2014; Bairlein, 2016; Murray et al., 2017), indicating that migratory species may be particularly sensitive to ongoing climatic and habitat changes and thus of particular concern for conservation (Wilcove & Wikelski, 2008).

The costs of long-range migration in insects have been little studied (Chapman et al., 2015). There is no fixed set of expectations of how movement behavior may shape species’ genetic diversity, as these may differ due to a variety of movements occurring in nature (Dingle, 1996, 2001; Dingle & Drake, 2007; Jonzén et al, 2011). The overall effective population size and the intensity of demographic fluctuations play large roles in the build-up and maintenance of genetic variation (Motro & Thomson, 1982; Whitlock, 1992). Migratory insects often exhibit vast population numbers (Chapman et al., 2011; Hu et al., 2016) and cyclic outbreaks (Wilson & Gatehouse, 1993; Chapuis et al., 2009; Satterfield et al., 2020). They also present some distinctive particularities in comparison with vertebrates that may shape demographic dynamics (Gao et al., 2020). These can include a higher reproductive potential, shorter generation times, or annual migrations that are completed through multiple generations. Migratory behavior may also counteract some of the factors that typically control insect population numbers, and result in comparatively larger population sizes in migratory compared to non-migratory species. First, the migratory strategy allows escape from deteriorating environmental conditions. As many migratory insects breed continuously without diapausing, different generations can exploit a succession of favorable breeding grounds that may reduce the probability for local bottlenecks. The high mobility associated with long-range migration may also lead to unusually large distribution ranges, which can increase the chances of finding optimal conditions. Finally, multigenerational migration allows insects to move into enemy-free space, most importantly escaping from parasitoids and predators that often increase in numbers in breeding areas as the season advances (Altizer et al., 2011; Bartel et al., 2011; Stefanescu et al., 2012; Chapman et al., 2015).

In contrast, theory states that gene flow will have a positive effect on the global level of genetic diversity, the effect being dependent on the migration rate and the level of diversity among immigrants (Leblois et al., 2004, 2006; Chapuis et al., 2009, 2014; Pfeiler & Markow, 2017). In migratory Lepidoptera, it is thus expected that seasonal migrations increase genetic diversity within migratory ranges due to throughout mixing between migratory groups, although ultimately depending on the degree of migratory connectivity between them (Webster et al., 2002; Gao et al., 2020). However, genetic diversity may also vary in non-migratory Lepidoptera due to other movement behaviours, which we summarise in Table 1. Since migratory and non-migratory species likely evolved under different selective pressures, we might expect that they will show distinctive patterns in the regularity and scale of their movements, and therefore in their population dynamics (Dingle, 2001; Webster et al., 2002; Dingle & Drake, 2007; Bhaumik & Kunte, 2020) (Table 1).

**Table 1.** Expected population-level outcomes from different types of movement in Lepidoptera potentially shaping connectivity and, ultimately, species’ genetic diversity.

|  | Sedentary | Dispersive<br>(other movements: e.g. ranging,<br>long-range foraging,<br>nomadism) | Migratory |
| --- | --- | --- | --- |
| **Nature of movement** | Resource-oriented,<br>not exploratory | Facultative response to<br>prevailing conditions,<br>exploratory, resource-oriented, terminated by<br>suitable resources | Obligate, anticipatory,<br>endogenously programmed<br>(pre-emption), temporary<br>inhibition of resource-<br>driven response |
| **Regularity** | Irregular | Irregularly recurrent<br>(between-year variability) | Seasonally recurrent<br>(constant) |
| **Proportion of the**<br>**population involved** | No displacement | Individual-scale: only<br>part of the population<br>may move | Population-scale:<br>whole population moves |
| **Distance between**<br>**individuals after movements** | Constant | Potentially increased | Nearly constant |
| **Extent of movements**<br>**(geographical scale)** | Short, within habitat | Short (between habitats)<br>to occasionally long-range (between regions),<br>usually one-way | Long-range, between<br>regions, usually two-way |
| **Duration of movements** | Very short | Short to long, but often<br>interrupted | Long and sustained |
| **Annual number of**<br>**generations involved** | All | None, one or more than one | All or nearly all |
| **Seasonal home-range**<br>**occupancy** | Always occupied<br>(not abandoned) | Always occupied<br>(not abandoned) | Temporarily<br>abandoned/colonized |
| **Safety of movement** | Predictable<br>(but not secured) | Unpredictable safe<br>habitat at destination | Predictable (but not<br>secured) safe habitat at<br>destination |
| **Predictability/regularity of**<br>**movement** | Yes | No | Yes |
| **Philopatry and migratory**<br>**connectivity** | High<br>(site-specific) | Very low | Low (region-specific) to<br>high (site-specific) |

Overall, we hypothesize that 1) a migratory life-style in Lepidoptera allows for a larger population size and, consequently, higher levels of genetic diversity in comparison to non-migratory species, and, 2) that demographic fluctuations between successive generations or seasons do not lead to severe bottlenecks or reduced genetic diversity at the species level. Here, we investigate whether migratory species of Lepidoptera show higher heterozygosity – a proxy for genetic diversity and effective population size – than non-migratory species. We estimate genomic heterozygosity across 381 individuals in 184 species in Lepidoptera, an order of insects whose life histories are reasonably well-known, and include a remarkable percentage of migratory species (ca. 3% found in butterflies; Chowdhury et al, 2021). As a paradigmatic case, we study in detail the demographic dynamics of the painted lady butterfly, *Vanessa cardui*. We generate whole-genome data and use coalescent modeling to infer population size and demographic dynamics across worldwide populations. This species is the most cosmopolitan of all butterflies (Shields, 1992), and performs annual latitudinal round-trip migrations involving 8-10 generations without diapausing (Talavera et al., 2018; Menchetti et al., 2019). Various studies have documented dramatic local fluctuations in *V. cardui*, both within and between years, seemingly coupled to natural climate oscillations (Williams, 1970; Pollard et al., 1998; Asher et al., 2001; Vandenbosch, 2003; Stefanescu et al., 2013; Menchetti et al., 2019; Hu et al., 2021; López-Mañas et al., 2022). We investigate whether these local fluctuations shape the demographic history of the species at a global scale, impacting the overall species genetic diversity, or alternatively, whether the migratory strategy of *V. cardui* reduces the risk of severe bottlenecks and thus leads to maintenance of high levels of genetic diversity.

## Materials and Methods

### Heterozygosity estimates

Heterozygosity, or heterozygosity rate, is the fraction of nucleotides within an individual that differ between the chromosomes they inherit from their parents, a basic genomic feature that can be computationally complex to estimate when genome assemblies are not available and sequence data is limited to unassembled reads (Bryc, 2013). We estimated levels of heterozygosity for 381 individuals belonging to 184 Lepidoptera species (Table S1, Figure S2) using GenomeScope v1.0 (Vurture et al., 2017), a method based on the statistical analysis of the *k-mer* profile (occurrence of substrings of length *k*) of raw short read sequencing data (Vurture et al., 2017). Homozygous genomes have a simple Poisson profile while heterozygous genomes have a characteristic bimodal profile (Kajitani et al., 2014). GenomeScope attempts in several rounds to measure genome-wide heterozygosity by fitting a mixed model that accounts for the distortion that sequencing errors, repeats and read duplications exert in these profiles (Vurture et al., 2017).

All species (including multiple individuals when available) with *>*1 Gb short reads available in the SRA archive in the NCBI database (accessed June 2021) were included in the analysis. A few individuals from laboratory lines were discarded to avoid biased heterozygosity estimates as a consequence of potential inbreeding. The selected species included taxa from five butterfly and eight moth families, and represented a diversity of movement behaviors (Table S1). Reads were right-trimmed to be of the same length for each sample and filtered for a minimum quality of 20 using BBDuk v38.61b (Bushnell, BBTools, 2020). We obtained coverage histograms with Jellyfish v2.2.6 (Marçais & Kingsford, 2011) using several *k-mer* lengths for each species (15, 17, 19, 21, 23, 25) to account for the variability in size and repetitiveness of the genomes. We provided these as input to GenomeScope to obtain genome-wide heterozygosity estimates and selected the best model for each species based on a minimum fit of 90%.

It is important to note that *k-mer* based heterozygosity estimates from a single sample are not comparable to absolute values of heterozygosity estimates obtained from SNP-based approaches involving multiple samples. As *k-mer* based heterozygosity is estimated using the number of mismatches between *k-mers*, this will be much closer to the estimates based on the number of segregating sites in the population, such as the Watterson estimator of genetic diversity (*θ*). Generally, estimates of heterozygosity based on GenomeScope might be slightly higher compared to those based on population estimates (Vurture et al., 2017).

### Behavioral scoring

We compiled evidence on migratory behavior from the literature for all taxa included in the dataset (Table S2). We systematically searched in Google Scholar and in a variety of regional field guidebooks for descriptors typically used in literature for Lepidoptera to describe movements of certain impact at geographical scale: the keywords “migration”, “migratory”, “migrating”, “dispersal”, “long-distance” and “long-range”, associated with each species name (including taxonomic synonyms). We also used as a working framework a list of nearly 600 butterfly species found to show evidence of migratory movements based on an exhaustive literature review (Chowdhury et al., 2021). All species of butterflies with migratory behavior retrieved from our literature search were also listed in Chowdhury et al (2021).

We scored the species in the dataset as belonging to one of three categories: sedentary, migratory and dispersive movements. Migratory behavior may be difficult to distinguish from other types of movement (Dingle, 1996; Dingle & Drake, 2007; Jonzén et al., 2011), and migration in Lepidoptera has generally been diagnosed based on population-level outcomes. For example, 92% of the butterfly species listed as migratory in the review by Chowdhury *et al*. (2021) were based only on field observations of directional mass movement, but only 8% of the species matched additional criteria at the experimental level. This 8% of species also had higher numbers of literature matches related to field-based observations of mass movements. As a general rule-of-thumb, we interpreted the absence of literature matches as absence of evidence of being a highly mobile species, and therefore scored these species as sedentary. Although sedentary (= resident) species may display round-trip movements within their home range in the search of resources (e.g. when foraging and/or exhibiting territorial behavior), these have a minor impact on the population level movement. In searching for literature matches, the information was carefully scrutinized for population-level evidence of migratory behavior and other types of movement (Dingle, 1996). Our rationale is summarized in Table 1.

Briefly, we define migratory species as those exhibiting stimuli-driven, endogenously programmed, and non-resource-oriented obligate movements outside their home range, involving seasonal shifts of whole populations. We define dispersal as the outcome of other types of movements that show a resource-oriented behavior or a facultative response to prevailing conditions involving exploration. Dispersal is generally unpredictable, leading to relevant geographical shifts, but ones where not all populations move. Our definition of dispersal may overlap, partially or completely, with the concept of ranging (sensu Dingle & Drake, 2007) or nomadism (sensu Jonzén et al, 2011). Generally, the information more commonly extracted from literature to distinguish migration from dispersal was related to the geographical scale of the movements (longer *vs* shorter ranges), the demographic scale (whole populations *vs* parts of the populations), the regularity (seasonal *vs* irregular recurrency) and the nature of the movement (obligate *vs* facultative). In most cases, species considered migratory were strongly supported by multiple publications and detailed ecology, while species considered dispersive showed evidence of important dispersal outcomes, but did not necessarily have consistent support from multiple sources, suggesting to us a better fit with species scored as dispersive rather than sedentary or migratory.

The only case where intraspecific behavioral variability was documented was in *Danaus plexippus*, where specimens from known migratory versus sedentary populations (Zhan et al., 2014; Freedman et al., 2020) were treated separately in downstream analyses. Overall, our behavioral categorization resulted in 16 species scored as migratory (17.64%), 69 as sedentary (65.66%), and 13 as dispersive (12.74%) (Figure 1, Table S1).

**Figure 1.**
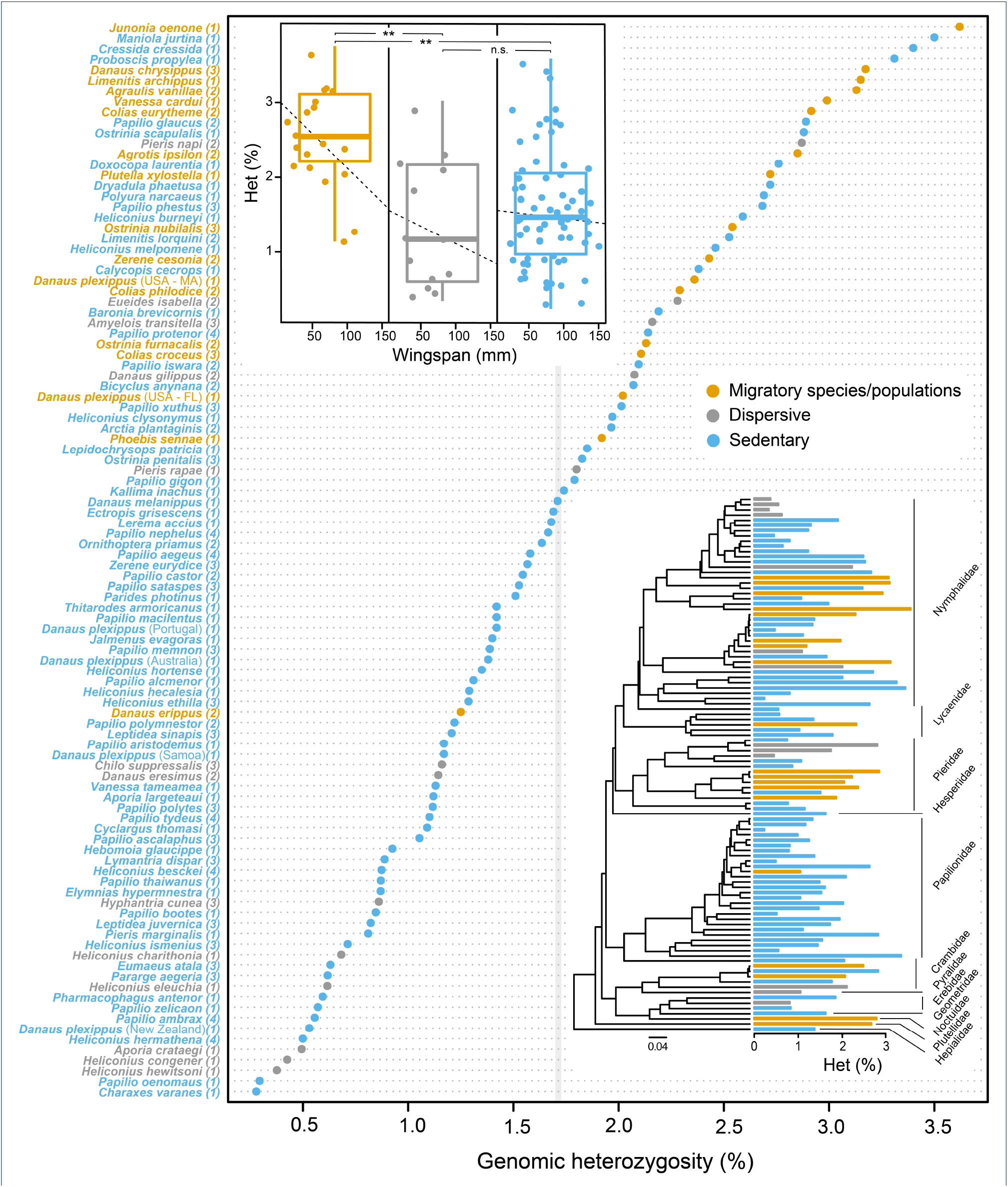
(a) Genomic heterozygosity for 185 individuals belonging to 97 Lepidoptera species calculated from short read Illumina reads using GenomeScope v1.0. Average estimates for species with several specimens are shown, with the exception of *Danaus plexippus*, for which migratory and non-migratory populations are illustrated. Numbers next to species names denote the number of individuals used for heterozygosity inference. The grey line indicates the average value for all analysed species. Species were classified as sedentary (in blue), migratory (in orange), and dispersive (in grey). (b) A pGLS analysis was used to test for significant deviation in heterozygosity between groups, accounting for wing size as proxy for body size (* = *P–value <* 0.05, ** = *P–value <* 0.01, *** = *P–value <* 0.001 and n.s. = non-signficant). (c) Phylogenetic diversity of the sampled species. Bars next to tips show average heterozygosity estimates.

### Comparative analyses

We built a phylogenetic framework including all species for which we previously inferred levels of genomic heterozygosity. We retrieved five of the most commonly used genes in Lepidoptera phylogenetics from GenBank (Table S3) and used BEAST2 (Bouckaert et al., 2014) to infer phylogenetic relationships. The substitution model was set to GTR+I+G with six gamma rate categories for each marker and a coalescent tree prior was set. We ran two independent MCMC chains of 100 million generations each, sampling values every 10,000 steps, and convergence was inspected in Tracer v.1.6.

We analysed the relationship between movement behaviour and heterozygosity levels using a phylogenetic generalized least squares (pGLS) regression to account for phylogenetic relatedness, as implemented in the R package ‘nlme’ (Pinheiro et al. 2016). For species where heterozygosity was estimated for multiple specimens, the mean values in the pGLS analysis were used. We also estimated the strength of the phylogenetic signal by inferring Pagel’s *λ* (Pagel, 1999) and Blomberg’s *K* (Blomberg, Garland & Ives 2003) in the model variables using the *phylosig* function in the R package ‘phytools’ (Revell, 2012) (Table S4).

Other life history traits could shape species’ heterozygosity levels and therefore act as confounding factors. As genetic diversity has been found to correlate with body size in butterflies (Mackintosh et al., 2019, Dapporto et al., 2019) and body size may correlate with population size (White et al., 2007), we included wingspan estimates obtained from the literature (Table S5) as a covariate in the pGLS test. In addition, because hybridization can elevate heterozygosity estimates, we excluded four potentially hybridizing taxa from our dataset (Table S1; Dupuis & Sperling, 2015, 2016; Herrell & Hazel, 1995; Mavárez et al., 2006).

### *Vanessa cardui* sampling and sequencing

We collected six *V. cardui* specimens from natural populations in five geographic regions: one specimen in Ethiopia (14U392; N 9° 23’ 54.096”, E 38° 49’ 25.323”; November 28, 2014), Namibia (15D327; N 17° 52’ 16.086”, E 19° 24’ 22.611”; April 21, 2015), Nepal (07F575; N 28° 25’ 0.635”, E 83° 49’ 2.114”; April 15, 2008), California, USA (15A205; N 38° 34’ 30.414”, O 121° 34’ 38.117’; March 24, 2015), and two specimens in Spain (15A708 and 15A710; N 40° 15’ 29.16”, O 1° 36’ 20.879”; May 22, 2015). Genomic DNA was extracted from the thorax using the Omniprep DNA extraction kit (G-Biosciences). We prepared two libraries for each specimen: a short-insert paired-end library (550 bp) from 2 g of DNA following TruSeq DNA PCR-Free LT sample preparation guide, and a mate pair library from 1 g of DNA following Nextera Mate Pair gel-free protocol. All libraries were sequenced at the genomic facilities of the University of Malaya (Malaysia) at 126 bp from both ends using Illumina HiSeq 2500 technology. The sequencing reads for all specimens have been deposited in the ENA (European Nucleotide Archive) database under accession numbers PRJEB43266.

### *Vanessa cardui* reference-based assemblies

Five re-sequenced samples, representatives from different geographic regions, were used for demographic inference. We obtained an average of 205 million reads from these samples (Table 2) and used BWA v0.7.12 (Li & Durbin, 2010) to map the reads to the available *V. cardui* genome (ilVanCard2.1, NCBI accession number GCA 905220365.1) (Lohse et al., 2022). An average of 88.6% of the reads were mapped and both mapping quality and mean depth of coverage across samples was high (Table 2), as assessed using Qualimap v2.2.1 (Okonechnikov, Conesa & García-Alcalde, 2016). Mapped sequences were sorted and indexed using SAMTOOLS v1.3 (Danecek et al., 2021). We also assessed the degree of completeness by analysis of the arthropod BUSCO v3.0.2 gene set (Simão et al., 2015) using gVolante (Nishimura, Hara & Kuraku, 2017). Of the 1013 genes queried, an average of 999 complete genes (98.63%) were retrieved from the assemblies, showing a high degree of completeness.

**Table 2.** Quality statistics of the five *V. cardui* genomes.

|  | Europe | N. America | S. Africa | E. Africa | Asia |
| --- | --- | --- | --- | --- | --- |
| Number of reads | 171,431,637 | 176,271,556 | 171,520,074 | 226,399,734 | 276,861,301 |
| Mapped reads | 160,199,554 | 163,200,113 | 122,289,727 | 208,310,949 | 259,227,222 |
| Percentage of mapped reads | 93.45 | 92.58 | 71.30 | 92.01 | 93.63 |
| Mean mapping quality | 48.60 | 48.32 | 48.19 | 48.29 | 47.92 |
| Mean coverage | 42.36 | 43.05 | 32.34 | 55.09 | 68.42 |
| GC content (%) | 32.88 | 33.05 | 33.17 | 32.72 | 33.41 |
| Heterozygosity (%) | 3.13 | 2.96 | 2.99 | 2.78 | 2.77 |
| BUSCO complete genes (%) | 999 (98.62%) | 997 (98.42%) | 1001 (98.82%) | 999 (98.62%) | 1000 (98.72%) |
| BUSCO complete + partial genes (%) | 1004 (99.11%) | 1003 (99.01%) | 1006 (99.31%) | 1005 (99.21%) | 1005 (99.21%) |

### *Vanessa cardui* demographic inference

We used the Pairwise Sequentially Markovian Coalescent (PSMC) (Li & Durbin, 2011) to investigate the demographic history of *V. cardui* samples from diverse geographical origins. This method relies on coalescence to infer past recombination and mutation events along the diploid genome of a single individual. As an input for PSMC, we used each individual’s consensus sequence generated with BCFTOOLS (Li, 2011). Variant calling was performed using the SAMTOOLS command *mpileup* (Li & Durbin, 2009). Repeat-rich regions are prone to bias effective population size (*N*_*e*_) estimates due to uncertainty in the quality of the variant calls, their population-level dynamics and deviations from the neutral expectations (Patil & Vijay, 2021). Thus, we excluded repeats as well as coding sequences in order to take into account only neutral sites. We used BEDtools (Quinlan & Hall, 2010) to exclude these regions from the consensus sequence using the available *V. cardui* genome and repeat/transposable element annotation as a reference (GCA 905220365.1: Lohse et al., 2022; Shipilina et al., 2022). Following Nadachowska-Brzyska et al., (2016), we excluded sites for which the mapping coverage was *<* 1/3 of the average read depth to minimize erroneously called homozygous sites, and sites where read depth was higher than twice the average in order to minimize the impact of potential erroneously collapsed regions in the assembly. We also filtered out positions with *Phred* score *<* 20. The upper limit of the TMRCA was set to 5 and parameter space for time intervals was set to “4+30*2+4+6+10”, so that at least 10 recombination events occurred after the twentieth iteration (Li & Durbin, 2011). In order to get estimates of variance in effective population size (*N*_*e*_), we performed 100 PSMC runs by randomly sampling with replacement 5 Mb sequence segments obtained from the consensus genome sequence. Average generation time was set to 0.11 years, corresponding to 9 generations per year (Talavera & Vila, 2017). The mutation rate was set to 2.9 × 10^−9^ mutations per year, corresponding to the estimate from the closest relative butterfly species, *Heliconius melpomene* (Keightley et al., 2015).

## Results

### Heterozygosity rates in Lepidoptera

From the heterozygosity estimates performed for 183 Lepidoptera species (Table S1), we excluded 53 species for which the GenomeScope model failed to converge, and 33 species for which the minimum model fit was below 90%. Heterozygosity in the 97 remaining species ranged between 0.21 % in *Heliconius hermathena* and 3.62 % in *Junonia oenone* (Figure 1, Table S1). The average heterozygosity for the six specimens of *V. cardui* was 2.95 % [2.77 – 3.13 %] (Table S1). No significant correlation was found between heterozygosity values and *k-mer* coverage (Pearson’s correlation, *t* = 0.62039, *P* –*value* = 0.537), suggesting that *k-mer* coverage was not a factor influencing our heterozygosity estimates. For 48 species, we could infer heterozygosity values in more than one individual (19 species with 2, 21 species with 3, 7 species with 4, and *V. cardui* with 6 individuals). Average intra-specific standard deviation in heterozygosity was 0.22, the minimum was 0.001, and the maximum was 0.86, indicating that estimates based on single individuals were generally robust. Intra-specific deviation did not correlate with species’ average heterozygosity (Pearson’s correlation, *t* = 0.62039, *P* –*value* = 0.537) (Figure S1), suggesting a random distribution of this variability. Additionally, outliers showing highest intra-specific differences belonged to the genus *Papilio* (a group known to hybridize frequently) (Herrell & Hazel, 1995; Dupuis & Sperling, 2015, 2016) or to be used as laboratory model systems (e.g. *Agrotis ipsilon*), both potentially affecting individual heterozygosity estimates. However, we found no evidence of either hybridization or inbreeding in individuals of these taxa in the original studies, so none were excluded from the analyses.

Migratory species had significantly higher heterozygosity values compared to sedentary species (mean ± s.e.: Migratory: 2.48 ± 0.65, Sedentary: 1.55 ± 0.79; pGLS_Strategy_: *F-stat =* 5.5978, *P-value* = 0.0200; Tukey Contrasts: *P-value* = 0.00514), and species categorized as dispersive (mean ± s.e.: 1.30 ± 0.835; Tukey Contrasts: *P-value* = 0.00893) (Figure 1). No significant differences in heterozygosity were found between the categories sedentary and dispersive (Tukey Contrasts: *P-value* = 0.08911). Wingspan had no significant effect on heterozygosity as a fixed effect (pGLS_Wingspan_: *F-stat =* 2.18108, *P-value* = 0.1429), although the interaction term was significant: wingspan and heterozygosity had a significant negative relationship in migratory species (pGLS_Wingspan*Strategy_: *F-stat =* 28.672282, *P-value <* 0.0001). Wingspan was independent from the explanatory variable ‘movement strategy’ (ANOVA, *F-stat* = 7.998896, *P-value* = 0.0057).

Heterozygosity estimates were phylogenetically independent as showed by low Blomberg’s *K* and Pagel’s *λ*, and migratory strategy showed weak phylogenetic signal, with *λ* and *K* values *<* 0.5 (Table S4). However, wingspan was distributed as expected under the Brownian motion (BM) model of trait evolution and exhibited strong phylogenetic dependence (*λ* = 0.99 and *K* = 1.50).

### *Vanessa cardui* demographic history

The estimated change in effective population size (*N*_*e*_) over time for five *V. cardui* individuals from the different sampled geographic regions is shown in Figure 2. The demographic histories of distant populations shared 1) a lack of abrupt *N*_*e*_ fluctuations and 2) a progressive population expansion that reaches a peak approximately ten thousand years ago (10 kya). We identify three phases in the demographic history of the different populations:

**Figure 2.**
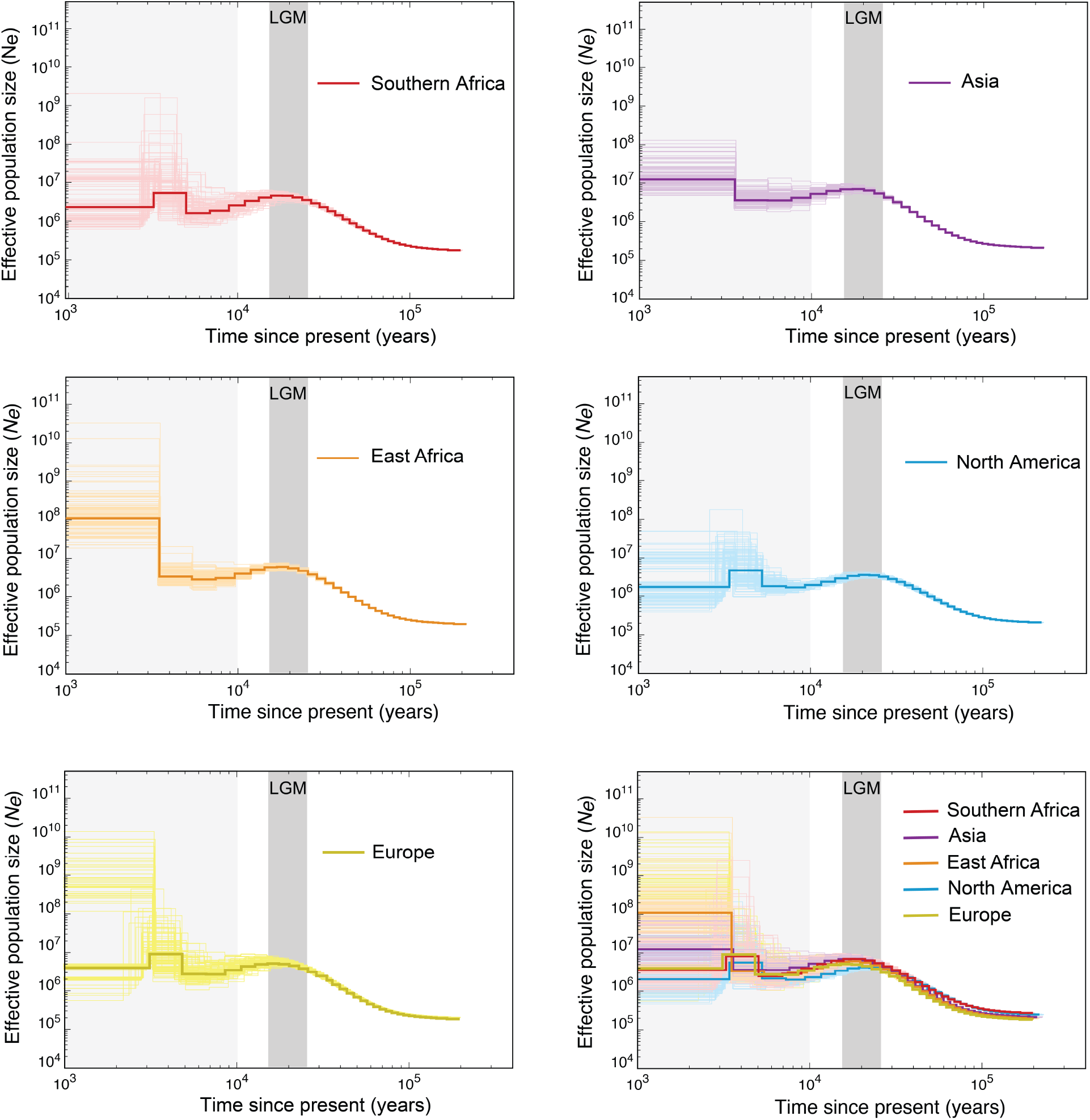
Demographic inference of *Vanessa cardui* populations from whole-genome data (excluding repeats and coding sequences). Pallid colored curves indicate PSMC estimates for 100 bootstrapped sequences for generation time g = 0.15 and mutation rate μ=0.3×10-8. The light gray shaded areas indicate the time range out of confidence of the PSMC estimations and the dark shaded gray indicate the Last Glacial Maximum (LGM) period. The estimated effective population size is shown in the y-axis, and coalescent time is shown in the x-axis. The low-right panel shows the different populations with PSMC trajectories superimposed for comparison.

1) 150 – 60 kya: a coalescent time for all populations at approximately 150 kya. Ancestral effective population size estimates for the species ranged from 10,000 to 15,000 individuals.

2) 80 – 20 kya: a period of slow but steady increase in population size, reaching *N*_*e*_ values between 200,000 and 800,000 individuals. The trajectories and the bootstrap replicates remained overlapping among populations.

3) 20 – 10 kya: a period of stabilization and slight decrease in *N*_*e*_, progressively differing among populations, suggesting an increase in continental-level population structure along with differences in *N*_*e*_. Estimates of *N*_*e*_ for this period range from 10^6^ to 10^7^ effective individuals.

## Discussion

### Genomic heterozygosity, effective population size and migration

We show that migratory Lepidoptera have higher heterozygosity values than non-migratory species. The highest levels of heterozygosity are found in species that are extremely abundant and/or are habitat generalists, many of them ranked as migratory or dispersive. *Junonia oenone* (heterozygosity = 3.62 %), for example, is migratory and one of the most abundant and widespread butterflies in Africa, flying year-round (including during the dry season), and thus avoiding diapause. Similar cases include the butterfly species *Danaus chrysippus* (3.23 %), *Colias eurytheme* (3.04 %), *Pieris napi* (2.76 %) and the moth species *Agrotis ipsilon* (3.34%), *Plutella xylostella* (2.72 %) and *Helicoverpa zea* (Anderson et al., 2018). Among the migratory species, *V. cardui* (2.95% ± 0.15) exhibits one of the highest levels of heterozygosity described in Lepidoptera (Li et al., 2019), ranking among the top 10 % of the most heterozygous species in our dataset (Figure 1). Other species found among the most heterozygous (Table S1) are suggested to hybridize, a process that can potentially be the cause of the observed high levels of heterozygosity. This concerns for example *Papilio polyxenes, P. joanae, P. machaon* (Herrell & Hazel, 1995; Dupuis & Sperling, 2015, 2016) and *Heliconius heurippa* (Mavárez et al., 2006).

A positive association between genetic diversity at neutral sites and *N*_*e*_ is expected, since larger populations have a higher input of novel mutations and reduced genetic drift (Wright, 1931; Kimura, 1983; Nevo, 1978; Leffler et al., 2012; Mackintosh et al., 2019). Our results suggest that migratory butterflies tend to have comparatively large *N*_*e*_. For example, in migratory monarch populations, *N*_*e*_ is inferred to be in the order of 10^7^ (Zhan et al., 2014; Talla et al., 2020), compared to *N*_*e*_ of 10^5^ – 10^6^ in sedentary populations (Zhan et al., 2014), the latter falling within a general range estimated for butterflies (Mackintosh et al., 2019). Our heterozygosity estimates for the monarch butterfly also followed a pattern of nearly twice the difference between behavioral categories: ca. 2 % in migratory and 1 ± 0.5 % in sedentary populations. For *V. cardui, N*_*e*_ were in the range of 10^6^ to 10^7^ at 10 kya. As census population size is expected to be larger than *N*_*e*_ (Fisher & Ford, 1947; Wright, 1948, Palstra & Ruzzante, 2008), one could debate whether such values are realistic. Several arguments support the existence of extraordinarily large population sizes in *V. cardui*. First, extremely large distribution ranges, typical of migratory insects, should facilitate elevated population sizes. The extreme dispersal capacity of *V. cardui* individuals is reflected in the vast migratory ranges that are covered annually across different continents. The species is also a polyphagous herbivore, and the diversity of larval hostplants and their phenology most likely expand opportunities for breeding success. As a typical r-strategist migratory insect, large population sizes may be advantageous in exploiting ephemeral flush vegetation, as well as in balancing the cost of higher mortality rates associated with migration (Dingle, 1978; Hill, 1983; Chapman et al., 2015). Moreover, the species has high reproductive potential (varying over several orders-of-magnitude in comparison to vertebrates), where single females may lay *>* 1,000 eggs (Benyamini, 2017), and can eventually behave as pests in particular breeding sites. Lastly, migration also facilitates higher population sizes by allowing escape from high parasitoid loads, an important factor exerting population control in insects (Jeffries & Lawton, 1984; Owen, 1987; Stefanescu et al., 2012; Chapman et al., 2015).

Although it is expected that highly abundant migratory species show high levels of heterozygosity given their greater opportunity to randomly share alleles between large populations, not all migratory species fall within this expectation. Indeed, there seem to be a few outliers that are not consistent with a correlation between genetic diversity and long-term population size. In Mackintosh et al., 2019, estimates of neutral genetic diversity for some migratory butterflies were among the bottom 15 % of European species, such as *Vanessa atalanta, Pontia daplidice* or *Pieris brassicae*. In our dataset, an exception is *Danaus erippus* (1.16 %), which appears to have a heterozygosity level below average. A deviation from a strict linear relationship between *N*_*e*_ and heterozygosity is not unexpected given a set of possible confounding factors. For example, recent geographic expansions and corresponding increases in population size, could lead to populations that have not yet reached mutation-drift equilibrium (Buffalo, 2021) – this has been suggested for example in *P. brassicae* and *P. rapae* (Mackintosh et al., 2019; Ryan et al., 2019). Another potential factor concerns the role that natural selection can play in decreasing neutral variation at physically linked sites (Ellegren & Galtier, 2016). In species with large population sizes, such ‘linked selection’ should be a more pervasive force since the efficiency of selection scales with *N*_*e*_ (Leffler et al., 2012; Corbett-Detig et al., 2015, Mackintosh et al., 2019; Talla et al., 2019). A paradigmatic case is the extinction of the passenger pigeon, once the most abundant migratory bird in North America and possibly in the world. Its surprisingly low genetic diversity has been suggested to be a collateral effect of natural selection (Murray et al., 2017). However, the effect of linked selection is also dependent on the recombination landscape, and detailed recombination maps will be needed to gain further insight into the relative effects of selection, drift and demography driving differences in heterozygosity between species with similar life histories. In *V. cardui*, PSMC results show no evidence that *N*_*e*_ drops despite the large population sizes that could enhance the action of selection, suggesting that selection has not been a major factor shaping the demography of this species.

Other potential factors shaping genomic heterozygosity involve migratory connectivity. Some species may have strong philopatry, resulting in high migratory connectivity between departure and destination, and thus little mixing between populations or migratory ranges (Battey et al., 2017; Gao et al., 2020), despite their long migratory movements. In the monarch butterfly migratory range, two (eastern and western) breeding and overwintering areas are repeatedly visited every year, and populations maintain their own phenotypic characteristics (Vandenbosch, 2003; Talla et al., 2020). This contrasts with the strategy of *V. cardui*, where individuals may be more opportunistic in their breeding destinations within each of the predictable suitable areas occupied in each multigenerational step (*i*.*e*. lower philopatry, in the sense that the populations do not rely on specific sites but rather on large regions). This results in lower migratory connectivity and thus higher allele sharing between different breeding sites.

Lastly, diapause may also influence migratory species’ genetic diversity and population size, as the number of generations per year is higher in species that breed continuously year-round compared to diapausing species. For example, the number of generations in *V. cardui* is twice that of *D. plexippus*, but also each *V. cardui* generation migrates and breeds in a new region, unlike the monarchs that keep three breeding generations within the same range (Flockhart et al., 2013). This uninterrupted movement and the high number of breeding regions/generations per year dramatically increase the possibilities of gene flow and is probably a relevant factor in explaining the very high levels of genomic heterozygosity in non-diapausing insects such as *V. cardui* and *J. oenone*.

The previously mentioned factors are expected to play a major role for migratory species, but these may not be observed in species that carry out other types of movements resulting in long-range dispersal. When dispersal is not seasonally recurrent and occurs at the individual level and not at the whole population scale, movement can increase the spatial distance between individuals, and eventually lower densities. While population admixture is expected to be large within the home range of these species, physical separation of individuals outside its range should limit the exchange of alleles and, eventually, may produce variation that is geographically structured. Species that can occasionally perform long-range dispersal have the potential to have larger distributional ranges than sedentary species, but not necessarily higher population size and heterozygosity, as our results suggest.

### Short-term migratory cost but long-term demographic stability

*Vanessa cardui* demographic history shows a stable population growth over time, high effective population sizes, and no apparent dramatic fluctuations or bottlenecks overall. This is consistent across populations worldwide. Long-term stability contrasts with the observation of frequent short-term population fluctuations and regional outbreaks observed in migratory insects between successive generations and years (Vanderbosch, 2003; Chapuis et al., 2009; Chapman et al., 2011; Hu et al., 2016; López-Mañas et al., 2022). Chapuis et al., (2009) found that outbreaking periods in the migratory locust (*Locusta migratoria*) did not strongly affect genetic diversity over time, either in outbreaking or non-outbreaking populations. Theory predicts that long-term effective size in populations recurrently fluctuating in their size can be approximated by calculating the harmonic mean of population size over all point estimates during times of fluctuations (Vucetich et al., 1997; Wang, 2005). The reason for this is that the loss of genetic diversity during periods of comparatively small *N*_*e*_ is expected to be higher than the increase in genetic diversity (due to mutational input) during periods of comparatively large *N*_*e*_. This means that the long-term *N*_*e*_ should be closer to the remission periods rather than to the outbreaking periods (Motro & Thomson 1982; Chapuis et al., 2009). Under this scenario, an outbreak remission should not be considered a bottleneck because it affects census but not *N*_*e*_. This rule probably applies to *V. cardui* and other migratory Lepidoptera, where regional fluctuations in census size may hardly affect the overall *N*_*e*_ of the species. Actually, outbreak events may lead to a homogenizing effect in the population structure of migratory insects (Chapuis et al., 2009; Anderson et al., 2018; Kaňuch et al., 2020). Given the large breeding ranges in most migratory insects, it is unlikely that local or even regional outbreaks or bottlenecks affect most populations across the whole range synchronically. Thus, a dilution or compensation effect of local demographic fluctuations takes place at the species level because of the extremely wide range of populations (Figure 3). For example, in *V. cardui* populations apparently cover entire continents or even intercontinental systems, such as the Palearctic-Africa system, and outbreaks may occur occasionally anywhere within large monthly breeding ranges (López-Mañas et al., 2022). This scenario is probably different in philopatric migratory vertebrates though, where breeding or overwintering ranges are smaller, more delimited, and thus more vulnerable to suffer bottlenecks and population declines if their habitats are disturbed. In this case, whole populations could be affected and genetic diversity loss is likely, with further population size recovery relying on the expansion of remaining populations (Figure 3).

**Figure 3.**
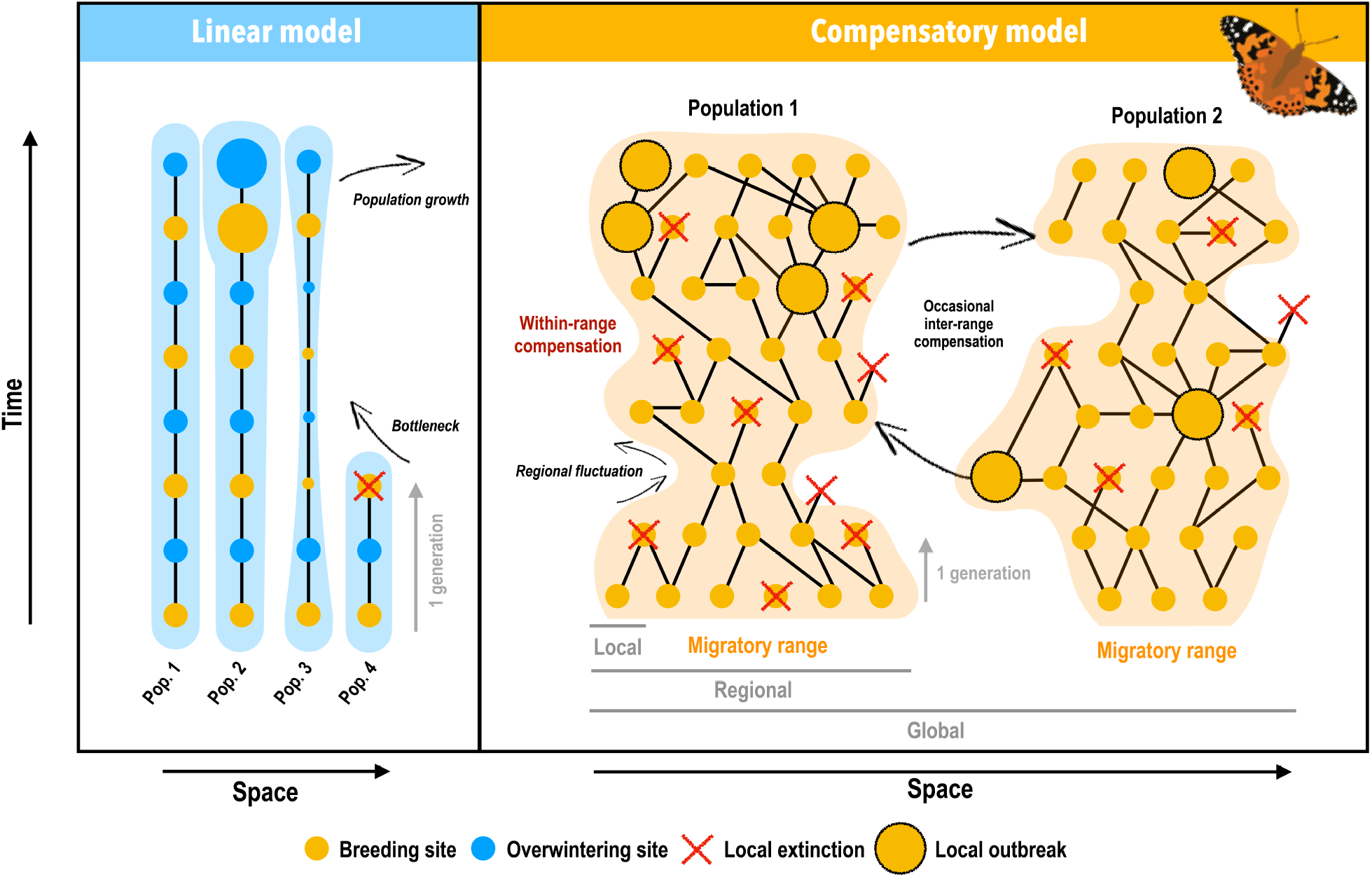
Conceptual models representing alternative population dynamics leading to different demographic histories in migratory animals. (a) A linear model, largely applicable to migratory vertebrates is characterized by long generation time, behaviorally-partitioned habitat (overwintering, stopovers and breeding ranges), and strong philopatry (high migratory connectivity) leading to little mixing between breeding sites. Population decreases or extinction events lead to bottlenecks and consequent loss of overall genetic diversity, while population growth occurs at population level. This model is likely to lead to fluctuations in long-term demographic history. (b) A compensatory model, generally distinctive to insects, where local/regional bottlenecks are diluted and compensated for by success in other areas. Large distributional ranges and breeding sites along them, low philopatry (low migratory connectivity), high reproductive potential (likely varying several orders-of-magnitude in a single generation in comparison to vertebrates), and short generation time are the main factors supporting this model. Long-term demographic history in this model is likely to show stability due to compensation within and occasionally between migratory ranges. The shaded areas represent the concept of migratory range, where individuals from a unique population circulate. The figure represents the spatial scale on the horizontal axis, from local breeding sites, to regional and global, and the temporal scale on the vertical axis. Dots represents breeding or overwintering sites, while crosses represent hypothetical extinction. The figure illustrates two extreme scenarios for migratory strategies in animals, assuming a wide variety of alternatives in between these two extremes. The case of *V. cardui* approximates well to the compensatory model.

Thus, a “compensatory” demographic model of migration in insects may result in a potential evolutionary advantage over the long-term, provided population sizes remain large enough compared to 1) sedentary species, where populations have limited ranges, thus making them potentially vulnerable to local habitat disturbance, unpredictable climatic events or parasitoid control, 2) strongly philopatric species, where success in different parts of the cycle rely on resources in specific sites, or 3) diapausing migratory species, where eventual bottlenecks could not be avoided during overwintering if important gatherings occur at a few key spots that may be exposed to environmental pressure.

### Delimiting populations in migratory insect species

Migratory ranges (or circuits) in insects could be defined as geographical ranges, even if parts of the ranges are only used temporarily, where gene flow is higher than between ranges. Migration in *V. cardui* follows a recurrent latitudinal progression that delineate a set of predictable multigenerational steps (Stefanescu et al., 2013; Menchetti et al., 2019), although gene flow may occur longitudinally to some extent. Thus, migratory ranges in this species might be generally defined as large longitudinal blocks where latitudinal gene flow is higher than longitudinal gene flow. However, the global configuration of these migratory ranges in *V. cardui* is not yet well understood. The species shares similar worldwide patterns of *N*_*e*_ variation over time, despite slight differences in North American and Southern African populations. This probably explains why present-day geographical populations share a common demographic history, implying that *V. cardui* is nearly panmictic, as found in other highly mobile species (Lyons et al., 2012; Raymond et al., 2013; Cristofari et al., 2016; Pfeiler and Markow, 2017; Talla et al., 2020), or at least that populations may circulate within extremely large distributional ranges. Taken together, our findings highlight the need to revisit the use of the population concept in *V. cardui* and in migratory insects in general. Dedicated population genetic studies are needed to infer historical and current patterns of gene flow and properly delimit migratory ranges.

## Acknowledgements

We thank Krystyna Nadachowska-Brzyska and Leonardo Dapporto for helpful methodological advice. This work was funded by the National Geographic Society (grant WW1-300R-18) and by the grant PID2020-117739GA-I00 from MCIN / AEI / 10.13039/501100011033 to G.T., by fellowship FPU19/01593 to A.G.-B., by grants from the Putnam Expeditionary Fund of the Museum of Comparative Zoology to G.T. and N.E.P., by projects PID2019-107078GB-I00 / MCIN / AEI / 10.13039/501100011033 and 2017-SGR-991 (Generalitat de Catalunya) to R.V. and G.T., by the University of Malaya (grant H50001-A-000027) to K.G.C., by the Swedish Collegium for Advanced Science (SCAS; Natural Sciences Programme, Knut and Alice Wallenberg Foundation, Postdoc funding) to D.S., by NSF grant DEB-1541560 to N.E.P., and by the Swedish Research Council FORMAS (grant 2019-00670) to N.B., G.T., and R.V. The authors also acknowledge support from the National Genomics Infrastructure in Stockholm funded by Science for Life Laboratory, the Knut and Alice Wallenberg Foundation and the Swedish Research Council, and SNIC/Uppsala Multidisciplinary Center for Advanced Computational Science for assistance with massively parallel sequencing and access to the UPPMAX computational infrastructure.

## Availability of data

Sequencing reads have been deposited in the ENA (European Nucleotide Archive) database under accession number PRJEB43266.

## Authors’ contributions

A.G-B., R.V., N.B. and G.T. conceived the study. G.T. and R.V. collected samples. H.K.W. and G.T. carried out laboratory work. A.G-B., V.T. and G.T. analysed data. A.G-B. and G.T. wrote the first version of the manuscript. All authors contributed in interpreting results and edited and approved the final version of the manuscript.

