## Supplementary material for "Migratory behavior is positively associated with genetic diversity in butterflies": Figure S1; Table S2, Table S3; Table S4

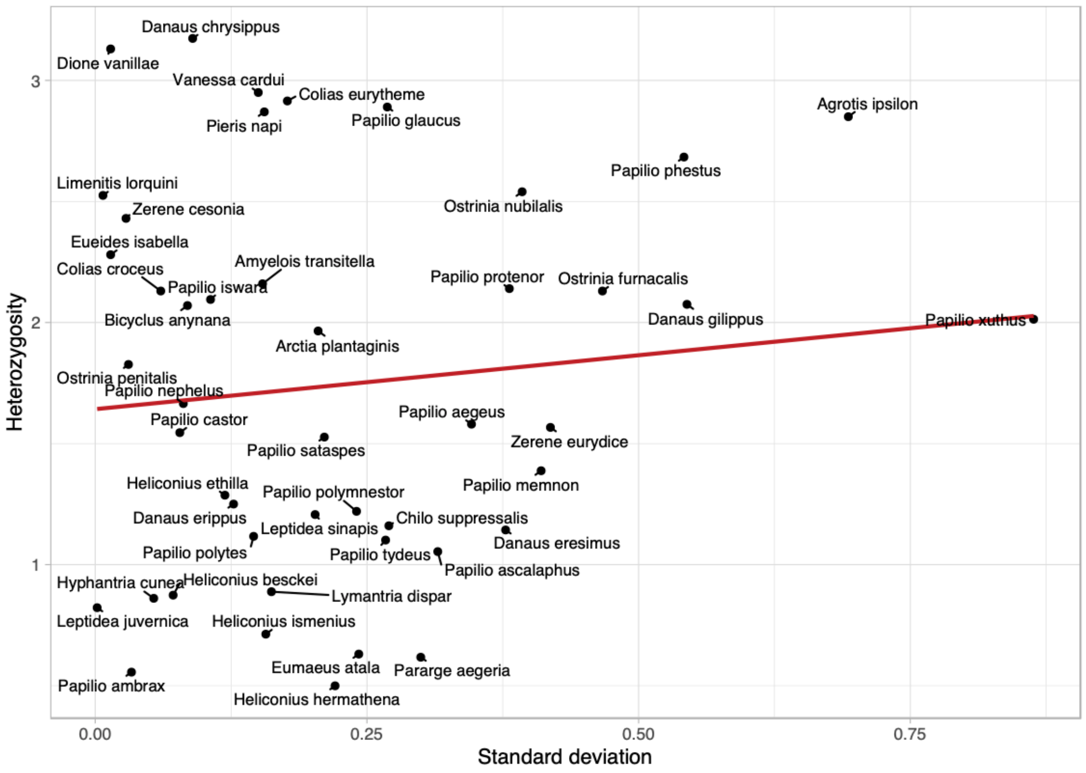


**Supplementary Figure S1.** Relationship between intra-specific average heterozygosity estimates (y-axis) obtained with GenomeScope and their standard deviations (x-axis). The average intra-specific standard deviation for 48 taxa accounting for ≥ 2 individuals was 0.22, the minimum was 0.001 and the maximum 0.86. The plot represents the distribution of this intra-specific variability as a function of their average heterozygosity values. There is no significant association between mean heterozygosity and standard deviation (Pearson's correlation, $t$ = 0.62039, *P–value* = 0.537), showing that intra-specific variability is not biased towards a specific level (high or low) of heterozygosity.

**Table S2.** Compiled evidence from literature associated to migratory behaviour for species included in heterozygosity estimates.

| Species | Strategy | References |
|---|---|---|
| *Agrotis ipsilon* | Migratory | Adden et al., 2020; Chang et al., 2019; Chang et al., 2018; Farrow & McDonald, 1987; Greenslade et al., 1999; Hendrix & Showers, 1992; Liu et al., 2015; Liu et al., 2016; Loublier et al., 1994; Loublier et al., 1993; Odiyo, 1975; Sappington & Showers, 1991, 1993; Sappington et al., 1995; Sappington et al., 1994; Sappington, 2018; Showers, 1997; Showers et al., 1993, Showers et al., 1986; Showers et al., 1989; Smelser et al., 1991; Sparks et al., 2007; Stancă-Moise, 2016; Thomas & William, 1992; Williams, 1930; Zeng et al., 2020 |
| *Amyelois transitella* | Dispersive | Andrews et al., 1980; Rovnyak et al., 2018; Sappington et al., 2014 |
| *Aporia crataegi* | Dispersive | Baker, 1969; Bolotov et al., 2013; Bolotov, 2021; Beshkov, 1996a; Jugovic et al., 2017; Larsen, 1977; Stancă-Moise, 2016; Williams, 1930 |
| *Aporia largeteaui* | Sedentary | No evidence found for long-distance migration. |
| *Arctia plantaginis* | Sedentary | No evidence found for long-distance migration. |
| *Baronia brevicornis* | Sedentary | No evidence found for long-distance migration. |
| *Byclus anyana* | Sedentary | No evidence found for long-distance migration. |
| *Calycopis cecrops* | Sedentary | No evidence found for long-distance migration. |
| *Charaxes varanes* | Sedentary | No evidence found for long-distance migration. |
| *Chilo suppressalis* | Dispersive | Pathak & Khan, 1994. |
| *Colias croceus* | Migratory | Baker, 1969; Bensusan et al., 2014; Haynes & Lavery 1990; John et al., 2006; Kettlewell, 1952; Lack, 1951; Larsen, 1975; Sparks et al., 2007; Stefanescu, 1997; Templado, 1976; Templado, 1977; Vos & Rutten, 1996; Williams, 1930 |
| *Colias eurytheme* | Migratory | David, 1982; Ferris, 1989, 1993; Fleishman, Murphy & Austin, 1999; Fisher, 1945; htttps://www.learnaboutbutterflies.com; Schweitzer, 2006; Williams, 1930; Shapiro, 1973, 1980a |
| *Colias philodice* | Migratory | Clarck, 1931; Fenton, 1919; Fisher, 1945; Williams, 1930 |
| *Cressida cressida* | Sedentary | No evidence found for long-distance migration. |
| *Cyclargus thomasi* | Sedentary | No evidence found for long-distance migration. |
| *Danaus chrysippus* | Migratory | Dingle et al., 1999; Gordon, 1984; Gugs et al., 2003; John et al., 2019; Koren et al., 2019; Larsen, 1975; Leigheb & Cameron-Curry, 1999; Lushai et al., 2003; Lushai et al., 2003; Samraoui, 1993; Samraoui & Benyacoub 1991; Singh et al., 2008; Smith et al., 1997; Smith et al., 1997; Williams, 1930. |
| *Danaus eresimus* | Dispersive | Benyamini, 2011; Calhoun, 1996; Srygley, 2001 |
| *Danaus erippus* | Migratory | Benyamini, 2011; Malcolm & Slager, 2015; Slager & Malcolm 2015; Williams, 1930 |
| *Danaus gilippus* | Dispersive | Hobson et al., 2021; Opler & Krizek, 1984; Saul-Gershenz et al., 2020; Williams, 1930 |
| *Danaus melanippus* | Sedentary | No evidence found for long-distance migration. |
| *Danaus plexippus (Australia)* | Dispersive | James & James, 2019; Dingle et al., 2019 |
| *Danaus plexippus (Florida)* | Migratory | Review in Oberhauser et al., 2015 |
| *Danaus plexippus (Hawaii)* | Sedentary | No evidence found for long-distance migration. |

| | | |
|---|---|---|
| *Danaus plexippus (Massachussets)* | Migratory | Review in Oberhauser et al., 2015 |
| *Danaus plexippus (New Zealand)* | Dispersive | James & James, 2019 |
| *Danaus plexippus (Portugal)* | Dispersive | Obregón et al, 2018 |
| *Danaus plexippus (Samoa)* | Sedentary | Recent colonization |
| *Doxocopa laurentia* | Sedentary | No evidence found for long-distance migration. |
| *Dryadula phaetusa* | Sedentary | No evidence found for long-distance migration. |
| *Ectropis grisescens* | Sedentary | No evidence found for long-distance migration. |
| *Elymnias hypermnestra* | Sedentary | No evidence found for long-distance migration. |
| *Eueides isabella* | Dispersive | Botello & Barrios, 2015 |
| *Eumaeus atala* | Sedentary | No evidence found for long-distance migration. |
| *Hebomoia glaucippe* | Sedentary | No evidence found for long-distance migration. |
| *Heliconius besckei* | Sedentary | No evidence found for long-distance migration. |
| *Heliconius burneyi* | Sedentary | No evidence found for long-distance migration. |
| *Heliconius charitonia* | Dispersive | Fleming et al., 2005; Kronforst & Fleming, 2001; Cardoso, et al., 2010 |
| *Heliconius clysonymus* | Sedentary | No evidence found for long-distance migration. |
| *Heliconius congener* | Sedentary | No evidence found for long-distance migration. |
| *Heliconius eleuchia* | Sedentary | No evidence found for long-distance migration. |
| *Heliconius ethilla* | Sedentary | No evidence found for long-distance migration. |
| *Heliconius hecalesia* | Sedentary | No evidence found for long-distance migration. |
| *Heliconius hermathena* | Sedentary | No evidence found for long-distance migration. |
| *Heliconius heurippa* | Sedentary | No evidence found for long-distance migration. |
| *Heliconius hewitsoni* | Sedentary | No evidence found for long-distance migration. |
| *Heliconius hortense* | Sedentary | No evidence found for long-distance migration. |
| *Heliconius ismenius* | Sedentary | No evidence found for long-distance migration. |
| *Heliconius melpomene* | Sedentary | No evidence found for long-distance migration. |
| *Hyphantria cunea* | Dispersive | Liu et al., 2006; Suzuki, 1981; Yamanaka, 2001 |
| *Jalmenus evagoras* | Sedentary | No evidence found for long-distance migration. |
| *Junonia oenone* | Dispersive | Timberlake & Childes, 2004 |
| *Kallima inachus* | Sedentary | No evidence found for long-distance migration. |
| *Lepidochrysops patricia* | Sedentary | No evidence found for long-distance migration. |
| *Leptidea juvernica* | Sedentary | No evidence found for long-distance migration. |
| *Leptidea sinapis* | Sedentary | No evidence found for long-distance migration. |
| *Lerema accius* | Sedentary | No evidence found for long-distance migration. |
| *Limenitis archippus* | Sedentary | No evidence found for long-distance migration. |
| *Limenitis lorquini* | Sedentary | No evidence found for long-distance migration. |
| *Lymantria dispar* | Sedentary | No evidence found for long-distance migration. |
| *Maniola jurtina* | Sedentary | No evidence found for long-distance migration. |
| *Ornithoptera priamus* | Sedentary | No evidence found for long-distance migration. |
| *Ostrinia furnacalis* | Migratory | Shen et al., 2020 |

| | | |
|---|---|---|
| *Ostrinia nubilalis* | Migratory | Sappington, T. W., 2018 |
| *Ostrinia penitalis* | Sedentary | No evidence found for long-distance migration. |
| *Ostrinia scapulalis* | Sedentary | No evidence found for long-distance migration. |
| *Papilio aegeus* | Sedentary | No evidence found for long-distance migration. |
| *Papilio alcmenor* | Sedentary | No evidence found for long-distance migration. |
| *Papilio ambrax* | Sedentary | No evidence found for long-distance migration. |
| *Papilio aristodemus* | Sedentary | No evidence found for long-distance migration. |
| *Papilio ascalaphus* | Sedentary | No evidence found for long-distance migration. |
| *Papilio bootes* | Sedentary | No evidence found for long-distance migration. |
| *Papilio castor* | Sedentary | No evidence found for long-distance migration. |
| *Papilio gigon* | Sedentary | No evidence found for long-distance migration. |
| *Papilio glaucus* | Sedentary | No evidence found for long-distance migration. |
| *Papilio iswara* | Sedentary | No evidence found for long-distance migration. |
| *Papilio joanae* | Sedentary | No evidence found for long-distance migration. |
| *Papilio machaon* | Dispersive | Sutton, 2009; Williams 1930 |
| *Papilio macilentus* | Sedentary | No evidence found for long-distance migration. |
| *Papilio memnon* | Sedentary | No evidence found for long-distance migration. |
| *Papilio nephelus* | Sedentary | No evidence found for long-distance migration. |
| *Papilio oenomaus* | Sedentary | No evidence found for long-distance migration. |
| *Papilio phestus* | Sedentary | No evidence found for long-distance migration. |
| *Papilio polymnestor* | Sedentary | No evidence found for long-distance migration. |
| *Papilio polytes* | Dispersive | Williams, 1930 |
| *Papilio polyxenes* | Sedentary | No evidence found for long-distance migration. |
| *Papilio protenor* | Sedentary | No evidence found for long-distance migration. |
| *Papilio sataspes* | Sedentary | No evidence found for long-distance migration. |
| *Papilio thaiwanus* | Sedentary | No evidence found for long-distance migration. |
| *Papilio tydeus* | Sedentary | No evidence found for long-distance migration. |
| *Papilio xuthus* | Dispersive | Williams, 1930 |
| *Papilio zelicaon* | Sedentary | No evidence found for long-distance migration. |
| *Pararge aegeria* | Sedentary | No evidence found for long-distance migration. |
| *Parides photinus* | Sedentary | No evidence found for long-distance migration. |
| *Pharmacophagus antenor* | Sedentary | No evidence found for long-distance migration. |
| *Phoebis sennae* | Migratory | Balciunas &Knopf, 1977; Chu, 2009; Dudley & Srygley, 2008; Hall et al. 2012; Hobson et al., 2021; Lenczewski, 1994; May 1992; Sprandel, 2001; Srygley, 2001; Srygley, 2001b; Srygley & Oliveira 2001; Urquhart & Urquhart, 1976; Walker & Littell, 1994; Waller, 1978, 1985, 1985b, 1991, 2001; Walker & Lenczewski, 1989; Walker & Riordan, 1981; Williams 1930 |
| *Pieris marginalis* | Sedentary | No evidence found for long-distance migration. |
| *Pieris napi* | Sedentary | No evidence found for long-distance migration. |
| *Pieris rapae* | Dispersive | Asher et al., 2001: Baker, 1969; Fric et al., 2006; Gilbert & Raworth, 2005; John et al., 2008; Jones et al., 1980; Larsen, 1975; Ohsaki et al., 1980; Ryan et al., 2019; Stefanescu et al., 2003; Walker, 2001; Williams, 1930 |

| | | |
|---|---|---|
| *Plutella xylostella* | Migratory | Campos et al., 2006; Chapman et al., 2002; Chen et al., 2020; Furlong et al., 2013; Harcourt, 1957; Shirai, 1995; Smith & Sears, 1982; Sparks et al., 2007; Williams, 1930; Yang et al., 2015; Yoshimatsu, 1991 |
| *Polyura narcaeus* | Sedentary | No evidence found for long-distance migration. |
| *Proboscis propylea* | Sedentary | No evidence found for long-distance migration. |
| *Thitarodes armoricanus* | Sedentary | No evidence found for long-distance migration. |
| *Vanessa cardui* | Migratory | Abbot, 1951; Giuliani & Shields, 1995; Lokki et al, 1978; Menchetti et al., 2019; Myres, 1985; Larsen 1975; Stefanescu et al., 2007; Stefanescu et al., 2012; Stefanescu et al., 2013; Stefanescu et al., 2016; Suchan et al., 2018; Talavera et al., 2018; Talavera & Vila, 2016; Tilden 1962; Vanderbosch, 2003; Williams, 1930; Williams, 1970. |
| *Vanessa tameamea* | Sedentary | No evidence found for long-distance migration/high dispersal behaviour. |
| *Zerene cesonia* | Migratory | Chu, 2007, 2009; Chowdhury et al., 2021; Hobson et al., 2021 |
| *Zerene eurydice* | Sedentary | No evidence found for long-distance migration. |

**Table S3.** List of GenBank codes used for phylogenetic inference.

| Species | COI | COII | EF | Wg | CAD |
|---|---|---|---|---|---|
| *Agraulis vanillae* | KF234586 | | | GQ864418 | GQ864600 |
| *Agrotis ipsilon* | KJ641991 | | | JQ786663 | JQ784293 |
| *Amyelois transitella* | HM388101 | | | | |
| *Aporia crataegi* | KU921257 | DQ463398 | KU921252 | EU141242 | EU141316 |
| *Aporia largeteaui* | KU921263 | KM669580 | | | |
| *Arctia plantaginis* | KX050271 | | | KX050658 | KX050194 |
| *Baronia brevicornis* | AF170866 | AF170866 | AF173406 | AY569044 | GQ864594 |
| *Bicyclus anyana* | HM241480 | HM241480 | HM241538 | | KR139361 |
| *Bombyx mandaria* | AB737944 | | | MH822604 | AB706327 |
| *Bombyx mori* | AB649195 | | EU490615 | EU141241 | EU141315 |
| *Calycopis cecrops* | HQ583417 | | | | |
| *Charaxes varanes* | GQ256872 | | GQ256991 | GQ256745 | |
| *Chilo suppressalis* | MK566582 | EU362118 | | JQ786745 | JQ784428 |
| *Colias croceus* | MN752706 | | MN752736 | | MN752805 |
| *Colias eurytheme* | AF044024 | AF044024 | FJ851634 | FJ851650 | EU032660 |
| *Colias philodice* | EU583855 | EU583882 | DQ157890 | AF537292 | KM046507 |
| *Cressida cressida* | AY919289 | | AY919294 | GQ268411 | |
| *Cyclargus thomasi* | HQ918891 | | | | |
| *Danaus chrysippus* | KP007634 | | KP007771 | KP007899 | |
| *Danaus eresimus* | GU659715 | | AY296135 | | |
| *Danaus erippus* | AY569158 | AY569158 | GU365931 | GU365966 | |
| *Danaus gilippus* | KP007668 | | KP007802 | KP007929 | |
| *Danaus melanippus* | KT286528 | | KT286221 | KT286048 | KT286348 |
| *Danaus plexippus (Australia)* | KF398175 | | AY296133 | | |
| *Danaus plexippus (Costa Rica)* | GU333884 | | | | |
| *Danaus plexippus (New Zealand)* | KF153721 | | | | |
| *Danaus plexippus (New Zealand)* | KF153721 | | | | |
| *Danaus plexippus (Ontario)* | KM550302 | | | AF233561 | EU141304 |
| *Danaus plexippus (Spain)* | KP871034 | | | | |

| | | | | | |
|---|---|---|---|---|---|
| *Depressaria pastinacella* | KT138815 | | | | |
| *Doxocopa laurentia* | MF547343 | | | | |
| *Dryadula phaetusa* | KF234610 | | | GQ864448 | KP073038 |
| *Ectropis grisescens* | KJ704363 | | | | |
| *Elymnias hypermnestra* | MW319394 | | | MW319473 | |
| *Eueides isabella* | AY748091 | AY748091 | | KF277432 | GQ864646 |
| *Eumaeus atala* | MW807685 | | | MT165253 | MT164668 |
| *Hebomoia glaucippe* | DQ082802 | | | KM046579 | KM046525 |
| *Heliconius besckei* | KP074764 | KP074764 | KP113968 | KP072822 | KP073052 |
| *Heliconius burneyi* | KP074765 | KP074765 | | KP072823 | KP073053 |
| *Heliconius charitonia* | KP074772 | KP074772 | KP113970 | KP072824 | KP073054 |
| *Heliconius clysonymus* | KP074773 | KP074773 | KP113971 | KP072826 | KP073056 |
| *Heliconius congener* | KP074775 | KP074775 | KP113973 | KP072828 | |
| *Heliconius eleuchia* | KP074788 | KP074788 | KP113988 | | KP073069 |
| *Heliconius ethilla* | KP074797 | KP074797 | KP114003 | KP072857 | KP073083 |
| *Heliconius hecalesia* | KP074800 | KP074800 | KP114010 | KP072860 | KP073086 |
| *Heliconius hermathena* | KP074804 | KP074804 | KP114015 | KP072864 | KP073089 |
| *Heliconius heurippa* | KP074807 | KP074807 | KP114018 | KP072865 | KP073090 |
| *Heliconius hewitsoni* | U08521 | U08521 | AY747926 | AF169881 | |
| *Heliconius hortense* | AY748043 | AY748043 | AY747929 | | |
| *Heliconius ismenius* | KP074815 | KP074815 | KP114023 | KP072870 | KP073096 |
| *Heliconius melpomene* | KP074817 | KP074817 | KP114024 | KP072871 | KP073097 |
| *Hyphantria cunea* | HM423646 | | | EU333645 | |
| *Jalmenus evagoras* | DQ456501 | DQ456501 | | KT286028 | KT286331 |
| *Junonia oenone* | AY788646 | | AY788765 | AY788525 | EU141332 |
| *Kallima inachus* | KX824673 | JN418279 | | KX824737 | |
| *Lepidochrysops patricia (dukei)* | GQ129024 | GQ129024 | | GQ128925 | |
| *Leptidea juvernica* | MW501464 | | | JF512971 | JF512764 |
| *Leptidea sinapis* | JF512602 | | | JF512984 | JF512748 |
| *Lerema accius* | KT598278 | KT598278 | | | |
| *Leucinodes orbonalis* | HQ991436 | | | | |
| *Limenitis archippus* | DQ205128 | DQ205128 | | EU433936 | |
| *Limenitis lorquini* | DQ205132 | DQ205132 | | GQ985321 | |
| *Lymantria dispar* | HM775744 | AB244663 | | EU333627 | |
| *Maniola jurtina* | KM020882 | | | AY090147 | EU141298 |
| *Ornithoptera priamus* | KT179873 | | KT179940 | KT180056 | |
| *Ostrinia furnacalis* | HQ991441 | AB121276 | | MK697104 | MK697190 |
| *Ostrinia nubilalis* | HM875015 | AB121311 | | MK697103 | MK459841 |
| *Ostrinia scapulalis* | KX041491 | EF622419 | | MK697102 | MK697188 |
| *Papilio aegeus* | KX557597 | KX557597 | KX558018 | KX558229 | |
| *Papilio alcmenor* | KX557677 | KX557677 | KX558096 | KX558308 | |
| *Papilio ambrax* | KX557624 | KX557624 | KX558045 | KX558256 | |
| *Papilio aristodemus* | KJ828883 | | KJ828926 | | |
| *Papilio ascalaphus* | KX557528 | KX557528 | KX557951 | KX558161 | |
| *Papilio bootes* | KX557587 | KX557587 | KX558008 | KX558218 | |
| *Papilio castor* | KX557641 | KX557641 | KX558061 | KX558273 | |
| *Papilio gigon* | KX557518 | KX557518 | KX557942 | KX558151 | |
| *Papilio glaucus* | AF044013 | AF044013 | AF044826 | GQ268406 | |
| *Papilio iswara* | KX557543 | KX557543 | KX557965 | KX558175 | |
| *Papilio joanae* | KJ363205 | KJ363205 | KJ363311 | | |
| *Papilio machaon* | AF044006 | AF044006 | AF044819 | AY569124 | |
| *Papilio macilentus* | KX557665 | KX557665 | KX558084 | KX558297 | |
| *Papilio memnon* | KX557698 | KX557698 | KX558117 | KX558329 | |
| *Papilio nephelus* | KX557699 | KX557699 | KX558118 | KX558330 | |
| *Papilio oenomaus* | KX557667 | KX557667 | KX558086 | KX558299 | |
| *Papilio phestus* | KX557553 | KX557553 | KX557975 | KX558185 | |
| *Papilio polymnestor* | KX557690 | KX557690 | KX558109 | KX558321 | |

| | | | | | |
|---|---|---|---|---|---|
| *Papilio polytes* | KX557695 | KX557695 | KX558114 | KX558326 | |
| *Papilio polyxenes* | AF044010 | AF044010 | AF044823 | | |
| *Papilio protenor* | KX557579 | KX557579 | KX558000 | KX558211 | |
| *Papilio sataspes* | KX557671 | KX557671 | KX557943 | KX558303 | |
| *Papilio thaiwanus* | AB377392 | AB377392 | JF681011 | | |
| *Papilio tydeus* | KX557545 | KX557545 | KX557967 | KX558177 | |
| *Papilio xuthus* | AF043999 | AF043999 | AF044838 | AY569123 | |
| *Papilio zelicaon* | AF044008 | AF044008 | AF044827 | | |
| *Pararge aegeria* | MH089836 | | | DQ338620 | EU141293 |
| *Parides photinus* | AF170877 | AF170877 | | DQ351127 | |
| *Pharmacophagus antenor* | AY919288 | AY919288 | AY919293 | GQ268410 | |
| *Phoebis sennae* | KM046828 | | AY870571 | | KM046547 |
| *Pieris macdunnoughi* | HQ978017 | | | | |
| *Pieris napi* | MN752730 | | MN752734 | MN752843 | MN752803 |
| *Pieris rapae* | GU372555 | | GU372646 | GQ283886 | GQ283572 |
| *Plutella xystosella* | NC_025322 | NC_025322 | GU829230 | | GU828102 |
| *Polyura narcaeus* | KU183740 (*athamas*) | AJ507641 | | | |
| *Proboscis propylea* | MN306904 | | | DQ338722 | |
| *Thitarodes armoricanus* | MK226958 | | | | |
| *Vanessa cardui* | HQ734908 | | HQ734947 | HQ734838 | HQ734873 |
| *Vanessa tameamea* | HQ734891 | | HQ734937 | HQ734829 | KJ648963 |
| *Zerene cesonia* | GU675579 | | | KM046597 | |
| *Zerene eurydice* | EU583851 | EU583878 | | | |

**Table S4.** Estimates of strength of phylogenetic signal Blomberg's *K* and Pagel's λ, inferred using *phylosig* function in the R package 'phytools'.

| Variable | Blomberg's K | | Pagel's λ | |
|---|---|---|---|---|
| | *K* value | ***P* –value** (based on 100000 randomizations) | *λ* value | ***P* –value** (based on LR test) |
| Heterozygosity | 0.106513 | 0.07461 | 7.24886e-05 | 1 |
| Strategy (Migratory/Sedentary/Dispersive) | 0.0682714 | 0.42399 | 0.455315 | 0.00445236 |
| Wingspan (Covariate) | 1.5041 | 1e-05 | 0.990525 | 2.42231e-28 |

### References (Table S2)


**Agraulis vanillae**

Arbogast, R. T. (1966). Migration of *Agraulis vanillae* (Lepidoptera, Nymphalidae) in Florida.

Beebe, W. (1950). Migration of Danaidae, Ithomiidae, Acraeidae and Heliconidae (butterflies) at Rancho Grande, north-central Venezuela. *Zoologica (New York)*, *35*, 57–68.

Lenczewski, B. (1994). Butterfly migration through the Florida peninsula.
O'Byrne, H. I. (1932). The migration and breeding of *Dione vanillae* in Missouri (Lepid.: Nymphalidae). *Entomol. News*, *43*, 97–99.
Opler, P. A. and G. O. Krizek (1984). Butterflies East of the Great Plains. Johns Hopkins University Press, Baltimore, MD. pp. 127–128.
Randolph, V. (1927). On the seasonal migrations of *Dione Vanillæ* in Kansas. *Annals of the Entomological Society of America*, *20*(2), 242–244.
Sharma, K. (2021). *Mechanisms of Lepidoptera edge effects and the impacts of temporal and environmental conditions on Gulf Fritillary Migration Surveys* (Doctoral dissertation, University of Georgia).
Sprandel, G. L. (2001). Fall dragonfly (Odonata) and butterfly (Lepidoptera) migration at St. Joseph Peninsula, Gulf County, Florida. *Florida Entomologist*, 234–238.
Srygley, R. B. (2001). Compensation for fluctuations in crosswind drift without stationary landmarks in butterflies migrating overseas. *Animal Behaviour*, *61*(1), 191–203.
Walker, T. J. (1980). Migrating Lepidoptera: are butterflies better than moths?. *The Florida Entomologist*, *63*(1), 79–98.
Walker, T. J. (1985). Butterfly migration in the boundary layer. *Migration: Mechanisms and adaptive significance*, 704–723.
Walker, T. J. (1991). Butterfly migration from and to peninsular Florida. *Ecological Entomology*, *16*(2), 241–252.
Walker, T. J. (2001). Butterfly migrations in Florida: seasonal patterns and long-term changes. *Environmental entomology*, *30*(6), 1052–1060.
Walker, T. J., & Littell, R. C. (1994). Orientation of Fall-migrating Butterflies in North Peninsular Florida and Source Areas. *Ethology*, *98*(1), 60–84.
Walker, T. J., & Riordan, A. J. (1981). Butterfly migration: are synoptic-scale wind systems important?. *Ecological Entomology*, *6*(4), 433–440.

### Agrotis ipsilon

Adden, A., Wibrand, S., Pfeiffer, K., Warrant, E., & Heinze, S. (2020). The brain of a nocturnal migratory insect, the Australian Bogong moth. *Journal of Comparative Neurology*, *528*(11), 1942–1963.
Chang, H., Guo, J. L., Fu, X. W., Hou, Y. M., & Wu, K. M. (2019). Orientation Behavior and Regulatory Gene Expression Profiles in Migratory *Agrotis ipsilon* (Lepidoptera: Noctuidae). *Journal of Insect Behavior*, *32*(1), 59–67.
Chang, H., Guo, J., Fu, X., Liu, Y., Wyckhuys, K. A., Hou, Y., & Wu, K. (2018). Molecular-assisted pollen grain analysis reveals spatiotemporal origin of long-distance migrants of a noctuid moth. *International journal of molecular sciences*, *19*(2), 567.
Farrow, R. A., & McDonald, G. (1987). Migration strategies and outbreaks of noctuid pests in Australia. *International Journal of Tropical Insect Science*, *8*(4), 531–542.
Greenslade, P., Farrow, R. A., & Smith, J. M. (1999). Long distance migration of insects to a subantarctic island. *Journal of Biogeography*, *26*(6), 1161–1167.
Hendrix, W. H., & Showers, W. B. (1992). Tracing black cutworm and armyworm (Lepidoptera: Noctuidae) northward migration using *Pithecellobium* and *Calliandra* pollen. *Environmental Entomology*, *21*(5), 1092–1096.
Liu, Y., Fu, X., Feng, H., Liu, Z., & Wu, K. (2015). Trans-regional migration of *Agrotis ipsilon* (Lepidoptera: Noctuidae) in north-east Asia. *Annals of the Entomological Society of America*, *108*(4), 519–527.
Liu, Y., Fu, X., Mao, L., Xing, Z., & Wu, K. (2016). Host plants identification for adult *Agrotis ipsilon*, a long-distance migratory insect. *International journal of molecular sciences*, *17*(6), 851.
Loublier, Y., Douault, P., Causse, R., Barthes, J., Bues, R., & Poitout, S. H. (1994). Utilisation des spectres polliniques recueillis sur *Agrotis* (Scotia) *ipsilon* Hufnagel (Noctuidae) comme indicateur des migrations. *Grana*, *33*(4-5), 276–281.

Loublier, Y., Douault, P., Causse, R., Barthes, J., Bues, R., & Poitout, S. H. (1993). Long-distance migration of black cutworm: pollen evidence (*Agrotis ipsilon*). *Apidologie (France)*.

Odiyo, P. O. (1975). *Seasonal distribution and migrations of* Agrotis ipsilon *(Hufnagel)(Lepidoptera, Noctuidae)* (No. 4). Centre for Overseas Pest Research (COPR).

Sappington, T. W., & Showers, W. B. (1991). Implications for migration of age-related variation in flight behavior of *Agrotis ipsilon* (Lepidoptera: Noctuidae). *Annals of the Entomological Society of America, 84*(5), 560–565.

Sappington, T. W., & Showers, W. B. (1993). Influence of larval starvation and adult diet on long-duration flight behavior of the migratory moth *Agrotis ipsilon* (Lepidoptera: Noctuidae). *Environmental entomology*, *22*(1), 141–148.

Sappington, T. W., Fescemyer, H. W., & Showers, W. B. (1995). Lipid and carbohydrate utilization during flight of the migratory moth, *Agrotis ipsilon* (Lepidoptera: Noctuidae). *Archives of Insect Biochemistry and Physiology*, *29*(4), 397–414.

Sappington, T. W., Showers, W. B., McNutt, J. J., Bernhardt, J. L., Goodenough, J. L., Keaster, A. J., ... & Way, M. O. (1994). Morphological correlates of migratory behavior in the black cutworm (Lepidoptera: Noctuidae). *Environmental entomology*, *23*(1), 58–67.

Sappington, T. W. (2018). Migratory flight of insect pests within a year-round distribution: European corn borer as a case study. *Journal of Integrative Agriculture*, *17*(7), 1485-1505.

Showers, W. B. (1997). Migratory ecology of the black cutworm. *Annual review of entomology*, *42*(1), 393–425.

Showers, W. B., Keaster, A. J., Raulston, J. R., Hendrix, W. H., Derrick, M. E., McCorcle, M. D., ... & Goodenough, J. L. (1993). Mechanism of southward migration of a noctuid moth [*Agrotis ipsilon* (Hufnagel)]: a complete migrant. *Ecology*, *74*(8), 2303–2314.

Showers, W. B., Keaster, A. J., Robinson, J. F., & Riley, T. J. (1986). Evidence of migration of the black cutworm adult into the US Corn Belt. *Long-range migrations of moths of agronomic importance to the United States and Canada: specific examples of occurrence and synoptic weather patterns conducive to migration: United States Department of Agriculture, Agri-cultural Research Service, ARS-43, Beltsville, Maryland, USA*, 10–24.

Showers, W. B., Whitford, F., Smelser, R. B., Keaster, A. J., Robinson, J. F., Lopez, J. D., & Taylor, S. E. (1989). Direct evidence for meteorologically driven long-range dispersal of an economically important moth. *Ecology*, *70*(4), 987–992.

Smelser, R. B., Showers, W. B., Shaw, R. H., & Taylor, E. S. (1991). Atmospheric trajectory analysis to project long-range migration of black cutworm (Lepidoptera: Noctuidae) adults. *Journal of economic entomology*, *84*(3), 879–885.

Sparks, T. H., Dennis, R. L., Croxton, P. J., & Cade, M. (2007). Increased migration of Lepidoptera linked to climate change. *European Journal of Entomology*, *104*(1), 139.

Stancă-Moise, C. (2016). Migratory species of butterflies in the surroundings of Sibiu (Romania). *Scientific Papers Series Management, Economic Engineering in Agriculture and Rural Development*, *16*(1), 319–324.

Thomas, W. S., & William, B. S. (1992). Reproductive maturity, mating status, and long-duration flight behavior of *Agrotis ipsilon* (Lepidoptera: Noctuidae) and the conceptual misuse of the oogenesis' flight syndrome by entomologists. *Environmental Entomology*, *21*(4), 677–688.

Williams, C. B. (1930). The migration of butterflies. *The Migration of Butterflies.*

Zeng, J., Liu, Y., Zhang, H., Liu, J., Jiang, Y., Wyckhuys, K. A., & Wu, K. (2020). Global warming modifies long-distance migration of an agricultural insect pest. *Journal of Pest Science*, *93*(2), 569–581.

**Amyelois transitella**

Andrews, K. L., Barnes, M. M., & Josserand, S. A. (1980). Dispersal and oviposition by navel orangeworm moths. *Environmental Entomology*, *9*(5), 525–529.

Rovnyak, A. M., Burks, C. S., Gassmann, A. J., & Sappington, T. W. (2018). Interrelation of mating, flight, and fecundity in navel orangeworm females. *Entomologia Experimentalis et Applicata*, *166*(4), 304–315.

Sappington, T. W., & Burks, C. S. (2014). Patterns of flight behavior and capacity of unmated navel orangeworm (Lepidoptera: Pyralidae) adults related to age, gender, and wing size. *Environmental entomology*, *43*(3), 696–705.

### Aporia crataegui

Baker, R. R. (1969). The evolution of the migratory habit in butterflies. *The Journal of Animal Ecology* 28, 703‑746.

Beshkov, S. (1996a). Migrant Lepidoptera species in Albania and Macedonia in 1995. *Atalanta* 27, 535‑543

Bolotov, I., Podbolotskaya, M., Kolosova, Y. S. & Zubrii, N. (2013). The current flow of migrants and its contribution to butterfly faunas (Lepidoptera, Rhopalocera) on marine islands with young allochthonous biota. *Biology Bulletin* 40, 78‑88.

Bolotov, I. N., Mizin, I. A., Zheludkova, A. A., Aksenova, O. V., Kolosova, Y. S., Potapov, G. S., ... & Gofarov, M. Y. (2021). Long-distance dispersal of migrant butterflies to the Arctic Ocean islands, with a record of Nymphalis xanthomelas at the northern edge of Novaya Zemlya (76.95 N). *Nota Lepidopterologica*, *44*, 73.

Jugovic, J., Črne, M., & Lužnik, M. (2017). Movement, demography and behaviour of a highly mobile species: A case study of the black-veined white, *Aporia crataegi* (Lepidoptera: Pieridae). *European Journal of Entomology*.

Larsen, T. B. (1977). Recent expansion in the range of *Aporia crataegi* (Linnaeus) in east Jordan (Pieridae). *Nota lepidopterologica*, *1*, 19–21.

Stancă-Moise, C. (2016). Migratory species of butterflies in the surroundings of Sibiu (Romania). *Scientific Papers Series Management, Economic Engineering in Agriculture and Rural Development*, *16*(1), 319–324.

### Chilo suppresalis

Pathak, M. D., & Khan, Z. R. (1994). Insect pests of rice. Manila, Philippines. International Rice Research Institute Newsletter 89pp. ISO 690

### Colias croceus

Baker, R. R. (1969). The evolution of the migratory habit in butterflies. *The Journal of Animal Ecology*, 703–746.

Bensusan, K. J., Nesbit, R., Perez, C. E., Tryjanowski, P., & Zduniak, P. (2014). Species composition and dynamics in abundance of migrant and sedentary butterflies (Lepidoptera) at Gibraltar during the spring period. *European Journal of Entomology*, *111*(4).

Haynes, R., & Lavery, T. (1990). Report on Migrant Insects in Ireland for 1989. *The Irish Naturalists' Journal, 23*(7), 277–279.

John, E., Coutsis, J. G., & Makris, C. (2006). A review of records for *Colias erate* (Esper,[1805])(Lep.: Papilionoidea Pieridae) in Cyprus: were they all yellow forms of *Colias croceus* (Geoffroy, 1785)?. *Entomologist's Gazette*, *57*, 3–12.

Kettlewell, H. B. D. (1952). A possible genetic explanation and understanding of migration of continuous brooded insects. *Nature*, *169*(4307), 832–833.

Lack, E. (1951). Migration of insects and birds through a Pyrenean pass. *The Journal of Animal Ecology*, 63–67.

Larsen, T. B. (1975). Provisional notes on migrant butterflies in Lebanon. *Atalanta Munnerstadt*, *62*, 62–74.

Sparks, T. H., Dennis, R. L., Croxton, P. J., & Cade, M. (2007). Increased migration of Lepidoptera linked to climate change. *European Journal of Entomology*, *104*(1), 139.

Stefanescu, C. (1997). Butterflies and moths (Insecta, Lepidoptera) recorded at sea off Eivissa and Barcelona (Western Mediterranean) in October 1996. *Bolletí de la Societat d'Història Natural de les Balears*, 51–56.

Templado, J. (1976). [An autumn migration of *Colias crocea* Geof.[Pieridae, butterfly] in Mandayona, Guadalajara [Spain]].[Spanish]. *Graellsia*.

Templado, J. (1977). A fall migration of *Colias crocea* Geof. in Mandayona, Guadalajara (Lepidoptera, Pieridae). *Graellsia*.

Vos, R. D., & Rutten, A. L. M. (1996). Migratory Lepidoptera in 1994 (fifty-fifth year). *Entomologische Berichten*, *56*(2), 17–27.

Williams, C. B. (1930). The migration of butterflies. *The Migration of Butterflies.*

### Colias eurytheme

David, G. (1982). The butterflies of Kent Island, Grand Manan, New Brunswick. *Journal of the Lepidopterists' Society* 36, 264-268.

Ferris, C. D. (1989). A new species of *Colias* from Utah (Pieridae: Coliadinae). *Bulletin of the Allyn Museum* 128,1-12.

Ferris, C. D. (1993). Reassessment of the *Colias* alexandra group, the Legume-feeding species, and preliminary cladistic analysis of the North American *Colias* (Pieridae: Coliadinae). *Bulletin of the Allyn Museum* 138,1-92.

Fisher, K. J. (1945). Two species of *Colias* migrating in central USA (Lep. Rhopalocera). *Proceedings of the Royal Entomological Society of London. Series A, General Entomology* *20*(10–12), 107–109.

Fleishman, E., Murphy, D. D. & Austin, G. T. (1999). Butterflies of the Toquima Range, Nevada: distribution, natural history, and comparison to the Toiyabe Range. *The Great Basin Naturalist 59*,50-62.

https://www.learnaboutbutterflies.com/North%20America%20-%20Colias%20eurytheme.htm

Schweitzer, D. F. (2006). The winter ecology of *Colias eurytheme* Boisduval (Pieridae) and its dependence on exotic legumes in Southern New Jersey. *Journal of the Lepidopterists' Society*, *60*(1), 51.

Williams, C. B. (1930). The migration of butterflies. *The Migration of Butterflies.*

Shapiro, A. M. (1973). Altitudinal migration of butterflies in the central Sierra Nevada. *Journal of Research on the Lepidoptera* 12, 231-235.

Shapiro, A. M. (1980a). Mediterranean climate and butterfly migration: an overview of the California fauna. *Atalanta* 11, 181-188.

### Colias philodice

Clark, A. H. (1931). Some observations on butterfly migrations. *The Scientific Monthly* 32, 150-155.

Fenton, C. L. (1919). Insect migration in floyd and adjoining counties of Iowa. American Midland Naturalist, 6(1), 13-15.

Fisher, K. J. (1945). Two species of *Colias* migrating in central USA (Lep. Rhopalocera). *Proceedings of the Royal Entomological Society of London. Series A, General Entomology* *20*(10–12), 107–109.

Williams, C. B. (1930). The migration of butterflies. *The Migration of Butterflies.*

### Danaus chrysippus

Dingle, H., Zalucki, M. P., & Rochester, W. A. (1999). Season-specific directional movement in migratory Australian butterflies. *Australian Journal of Entomology*, *38*(4), 323–329.

Gordon, I. J. (1984). Polymorphism of the tropical butterfly, *Danaus chrysippus* L., in Africa. *Heredity*, *53*(3), 583–593.

Gugs, L., Gordon, I.J., Smith D.A.S. (2003). Evidence from mitochondrial DNA supports earlier records of African queen butterflies (*Danaus chrysippus)* migrating in East Africa. *Journal of East African Natural History* 92, 119-125

John, E., Hardman, M., & Smith, M. (2019). How important are olfactory cues for host-plant detection by migrating *Danaus chrysippus* (Linnaeus, 1758)(Lepidoptera: Nymphalidae, Danainae) in Cyprus?. *Entomologist's Gazette*, *70*(4), 223–238.

Koren, T., Dender, D., Ilić, B., & Martinović, M. On the distribution and status of the African monarch (*Danaus chrysippus* (Linnaeus, 1758); Lepidoptera: Nymphalidae) in Croatia.
Larsen, T. B. (1975). Provisional notes on migrant butterflies in Lebanon. *Atalanta Munnerstadt*, *62*, 62–74.
Leigheb, G., & Cameron-Curry, V. (1999). Observations on the presence of Danaus chrysippus (Linné, 1758) in the Mediterranean area, with special reference to Italy (Lepidoptera, Danaidae). *Linneana Belgica (Belgium)*.
Lushai, G., Gordon, I. J., & Smith, D. A. (2003). Evidence from mitochondrial DNA supports earlier records of African queen butterflies (*Danaus chrysippus*) migrating in East Africa. *Journal of East African Natural History*, *92*(1), 119–125.
Lushai, G., Smith, D. A., Gordon, I. J., Goulson, D., Allen, J. A., & Maclean, N. (2003). Incomplete sexual isolation in sympatry between subspecies of the butterfly *Danaus chrysippus* (L.) and the creation of a hybrid zone. *Heredity*, *90*(3), 236–246.
Samraoui, B. (1993). Migration of the African monarch *Danaus chrysippus* (L.) and the African migrant Catopsilia florella (Fabr.) in Mauritania (Lepidoptera: Danaidae, Pieridae). *Nota Lepidopterologica*, *16*(1), 68–70.
Samraoui, B., & Benyacoub, S. (1991). A large migration of the plain tiger, *Danaus chrysippus* L., through northeastern Algeria (Lepidoptera: Danaidae). *Nota Lepidopterologia*, *14*, 99–100.
Singh, V. K., Joshi, P. C., & Joshi, B. D. (2018). Molecular data suggest population expansion and high level of gene flow in the Plain Tiger (*Danaus chrysippus*; Nymphalidae: Danainae). *Mitochondrial DNA Part B*, *3*(2), 707–712.
Smith, D. A., & Owen, D. F. (1997). Colour genes as markers for migratory activity: The butterfly *Danaus chrysippus* in Africa. *Oikos*, 127–135.
Smith, D. A., Owen, D. F., Gordon, I. J., & Lowis, N. K. (1997). The butterfly *Danaus chrysippus* (L.) in East Africa: polymorphism and morph-ratio clines within a complex, extensive and dynamic hybrid zone. *Zoological Journal of the Linnean Society*, *120*(1), 51–78.
Williams, C. B. (1930). The migration of butterflies. *The Migration of Butterflies.*

**Danaus eresimus**

Benyamini, D. (2011). Danaus eresimus (Cramer, 1777) new to Chile and the status in that country of Danaus erippus (Cramer, 1776). *Tropical Lepidoptera Research*, 98–99.
Calhoun, J. V. (1996). Conquering soldiers: the successful invasion of Florida by *Danaus eresimus* (Lepidoptera: Nymphalidae). *Holarctic Lepidoptera*, 7–18.
Srygley, R. B. (2001). Compensation for fluctuations in crosswind drift without stationary landmarks in butterflies migrating overseas. *Animal Behaviour*, *61*(1), 191–203.

**Danaus erippus**

Benyamini, D. (2011). *Danaus eresimus* (Cramer, 1777) new to Chile and the status in that country of *Danaus erippus* (Cramer, 1776). *Tropical Lepidoptera Research*, 98–99.
Malcolm, S. B., & Slager, B. H. (2015). Migration and host plant use by the southern monarch, Danaus erippus. *Monarchs in a Changing World: Biology and Conservation of an Iconic Butterfly*, 225.
Slager, B. H., & Malcolm, S. B. (2015). Evidence for partial migration in the southern monarch butterfly, Danaus erippus, in Bolivia and Argentina. *Biotropica*, *47*(3), 355–362.
Williams, C. B. (1930). The migration of butterflies. *The Migration of Butterflies.*

**Danaus gilippus**

Saul-Gershenz, L., Grodsky, S. M., & Hernandez, R. R. (2020). Ecology of the western queen butterfly Danaus gilippus thersippus (Lepidoptera: Nymphalidae) in the Mojave and Sonoran Deserts. *Insects*, *11*(5), 315.
Opler, P. A., & Krizek, G. O. (1984). *Butterflies east of the Great Plains: an illustrated natural history.* Johns Hopkins University Press.

Hobson, K. A., Kusack, J. W., & Mora-Alvarez, B. X. (2021). Origins of six species of butterflies migrating through northeastern Mexico: New insights from stable isotope (δ2H) analyses and a call for documenting butterfly migrations. *Diversity*, *13*(3), 102.

Williams, C. B. (1930). The migration of butterflies. *The Migration of Butterflies.*

#### Danaus plexippus

**Original range:**

Oberhauser, K. S., & Solensky, M. J. (Eds.). (2004). *Monarch butterfly biology & conservation*. Cornell university press.

Oberhauser, K. S., Nail, K. R., & Altizer, S. (Eds.). (2015). *Monarchs in a changing world: biology and conservation of an iconic butterfly*. Cornell University Press.

**Recent colonizations:**

Dingle, H., Zalucki, M. P., & Rochester, W. A. (1999). Season-specific directional movement in migratory Australian butterflies. *Australian Journal of Entomology*, *38*(4), 323–329.

James, D. G., & James, T. A. (2019). Migration and Overwintering in Australian Monarch Butterflies (*Danaus plexippus* (L.)(Lepidoptera: Nymphalidae): a Review with New Observations and Research Needs. *The Journal of the Lepidopterists' Society*, *73*(3), 177–190.

Obregón, R., Jordano, D., Cuadrado, M., Moreno-Benítez, J. M., Fernández Haeger, J. 2018. Dispersal of the monarch butterfly (*Danaus plexippus*) over southern Spain from its breeding grounds. *Animal Biodiversity and Conservation*.

#### Eueides Isabella

Botello, F. D., & Barrios, H. (2015). Migración invernal de mariposas en la playa el Agallito, Chitré, Provincia de Herrera, Panamá. *Scientia*, *25*(2), 15–33.

#### Heliconius charitonia

Fleming, T. H., Serrano, D., & Nassar, J. (2005). Dynamics of a subtropical population of the zebra longwing butterfly Heliconius charithonia (Nymphalidae). *Florida Entomologist*, 169–179.

Kronforst, M. R., & Fleming, T. H. (2001). Lack of genetic differentiation among widely spaced subpopulations of a butterfly with home range behaviour. *Heredity*, *86*(2), 243–250.

Cardoso, M. Z. (2010). Reconstructing seasonal range expansion of the tropical butterfly, Heliconius charithonia, into Texas using historical records. *Journal of Insect Science*, *10*(1), 69.

#### Hyphantria cunea

Liu, H., Luo, Y., Wen, J., Zhang, Z., Feng, J., & Tao, W. (2006). Pest risk assessment of *Dendroctonus valens*, *Hyphantria cunea* and *Apriona swainsoni* in Beijing. *Frontiers of Forestry in China*, *1*(3), 328–335.

Suzuki, N., & Kunimi, Y. (1981). Dispersal and survival rate of adult females of the fall webworm: *Hyphantria cunea* Drury (Lepidoptera: Arctiidae): Using the nuclear polyhedrosis virus as a marker. *Applied entomology and zoology*, *16*(4), 374–385.

Yamanaka, T., Tatsuki, S., & Shimada, M. (2001). Flight characteristics and dispersal patterns of fall webworm (Lepidoptera: Arctiidae) males. *Environmental entomology*, *30*(6), 1150–1157.

#### Junonia oenone

Timberlake, J. R., & Childes, S. L. (2004). Biodiversity of the Four Corners Area: Technical Reviews Volume Two (Chapter 10). *Occasional Publications in Biodiversity, 15*, 179.

#### Ostrinia nubilalis

Sappington, T. W. (2018). Migratory flight of insect pests within a year-round distribution: European corn borer as a case study. *Journal of Integrative Agriculture 17*(7): 1485–1505.

Showers, W. B. (1979). Effect of diapause on the migration of the European corn borer into the southeastern United States. In *Movement of highly mobile insects: concepts and methodology in research; proceedings of a conference, Movement of Selected Species of Lepidoptera in the Southeastern United States, Raleigh, North Carolina, April 9-11, 1979, edited by RL Rabb and GG Kennedy*. Raleigh, NC, University Graphics, 420–430.

**Ostinia furnacalis**

Shen, X., Fu, X., Huang, Y., Guo, J., Wu, Q., He, L., ... & Wu, K. (2020). Seasonal Migration Patterns of *Ostrinia furnacalis* (Lepidoptera: Crambidae) Across the Bohai Strait in Northern China. *Journal of economic entomology*, *113*(1), 194–202.

Shen, X., Guo, J., & Wu, K. (2021). Effects of Biotic and Abiotic Factors on Flight Performance of Ostrinia furnacalis. *Journal of Insect Behavior*, *34*(4), 240–253.

Shirai, Y. (1998). Laboratory evaluation of flight ability of the Oriental corn borer, Ostrinia furnacalis (Lepidoptera: Pyralidae). *Bulletin of entomological research*, *88*(3), 327–333.

**Papilio machaon**

Sutton, S. L. (2009). *South Caspian Insect Fauna, 1961. Transactions of the Royal Entomological Society of London, 118(3), 51–72.*

Williams, C. B. (1930). The migration of butterflies. *The Migration of Butterflies.*

**Phoebis sennae**

Balciunas, J., & Knopf, K. (1977). Orientation, flight speeds, and tracks of three species of migrating butterflies. *Florida Entomologist*, 37–39.

Chu, J. (2009). Inventories of Butterflies in Boulder County. *Unpublished report for Boulder County Parks and Open Space, Boulder, CO.*

Dudley, R., & Srygley, R. B. (2008). Airspeed adjustment and lipid reserves in migratory Neotropical butterflies. *Functional Ecology*, 264–270.

Hall, D. W., Minno, M. C., & Walker, T. J. (2012). Cloudless Sulphur *Phoebis sennae* (Linnaeus)(Insecta: Lepidoptera: Pieridae: Coliadinae). Entomology and Nematology Department, Florida Cooperative Extension Service, Institute of Food and Agricultural Sciences, University of Florida.

Hobson, K. A., Kusack, J. W., & Mora-Alvarez, B. X. (2021). Origins of six species of butterflies migrating through northeastern Mexico: New insights from stable isotope (δ2H) analyses and a call for documenting butterfly migrations. *Diversity*, *13*(3), 102.

Lenczewski, B. (1994). Butterfly migration through the Florida peninsula.

May, P. G. (1992). Flower selection and the dynamics of lipid reserve in two nectarivorous butterflies. *Ecology*, *73*(6), 2181–2191.

Sprandel, G. L. (2001). Fall dragonfly (odonata) and butterfly (lepidoptera) migration at St. Joseph Peninsula, Gulf County, Florida. *Florida Entomologist*, 234–238.

Srygley, R. B. (2001). Compensation for fluctuations in crosswind drift without stationary landmarks in butterflies migrating overseas. *Animal Behaviour*, *61*(1), 191–203.

Srygley, R. B. (2001). Sexual differences in tailwind drift compensation in *Phoebis sennae* butterflies (Lepidoptera: Pieridae) migrating over seas. *Behavioral Ecology*, *12*(5), 607–611.

Srygley, R. B., & Oliveira, E. G. (2001). Sun compass and wind drift compensation in migrating butterflies. *Journal of Navigation 54*(3), 405–417.

Urquhart, F. A., & Urquhart, N. R. (1976). Migration of butterflies, along the Gulf Coast of northern Florida. *Journal of the Lepidopterists' Society 30*, 59–61.

Walker T. J., Littell R. C. (1994). Orientation of fall migrating butterflies in north peninsular Florida and source areas. *Ethology, 98*(1): 60–84.

Walker, T. J. (1978). Migration and re-migration of butterflies through north peninsular Florida: quantification with Malaise traps. *Journal of the Lepidopterists' Society 32*(3), 178–190.

Walker, T. J. (1985). Butterfly migration in the boundary layer. *Migration: Mechanisms and adaptive significance*, 704–723.
Walker, T. J. (1985). Permanent traps for monitoring butterfly migration: Tests in Florida, 1979-84. *Journal of the Lepidopterists Society*, *39*(4), 313–320.
Walker, T. J. (1991). Butterfly migration from and to peninsular Florida. *Ecological Entomology*, *16*(2), 241–252.
Walker T. J. (2001). Butterfly migrations in Florida: seasonal patterns and long-term changes. *Environmental Entomology, 30(*6), 1052–1060.
Walker, T. J., & Lenczewski, B. (1989). An inexpensive portable trap for monitoring butterfly migration. *Journal of the Lepidopterists' Society*, *43*(4), 289–298.
Walker, T. J., & Riordan, A. J. (1981). Butterfly migration: are synoptic-scale wind systems important?. *Ecological Entomology*, *6*(4), 433–440.
Williams, C. B. (1930). The migration of butterflies. *The Migration of Butterflies.*

**Pieris napi**

Baker, R. R. (1969). The evolution of the migratory habit in butterflies. *The Journal of Animal Ecology*, 703-746.
Stefanescu, C., Peñuelas, J., & Filella, I. (2003). Effects of climatic change on the phenology of butterflies in the northwest Mediterranean Basin. *Global change biology*, *9*(10), 1494–1506.
Larsen, T. B. (1975). Provisional notes on migrant butterflies in Lebanon. *Atalanta Munnerstadt*, *62*, 62–74.

**Pieris rapae**

Asher, J., Warren, M., Fox, R., Harding, P., Jeffcoate, G., & Jeffcoate, S. (2001). *The millennium atlas of butterflies in Britain and Ireland*. Oxford University Press.
Baker, R. R. (1969). The evolution of the migratory habit in butterflies. *The Journal of Animal Ecology*, 703–746.
Fric, Z., Klimova, M., & Konvicka, M. (2006). Mechanical design indicates differences in mobility among butterfly generations. *Evolutionary Ecology Research*, *8*(8), 1511–1522.
Gilbert, N., & Raworth, D. A. (2005). Movement and migration patterns in *Pieris rapae* (Pieridae). *Journal of the Lepidopterists' Society 59*, 10–18.
John, E., Cottle, N., McArthur, A., & Makris, C. (2008). Eastern Mediterranean migrations of *Pieris rapae* (Linnaeus, 1758)(Lepidoptera: Pieridae): observations in Cyprus, 2001 and 2007. *Entomologists Gazette*, *59*(2), 71.
Jones, R. E., Gilbert, N., Guppy, M., & Nealis, V. (1980). Long-distance movement of *Pieris rapae*. *The Journal of Animal Ecology*, 629–642.
Larsen, T. B. (1975). Provisional notes on migrant butterflies in Lebanon. *Atalanta Munnerstadt*, *62*, 62–74.
Ohsaki, N (1980). Comparative population studies of three *Pieris* butterflies, *P. rapae, P. melete* and *P. napi,* living in the same area II. utilization of patchy habitats by adults through migratory and non-migratory movements. *Res. Popul. Ecol* 22, 163-183.
Ryan SF, Lombaert E, Espeset A, Vila R, Talavera G, Dinca V, Renshaw MA, Eng MW, Doellman MM, Hornett EA, Li Y, Pfrender ME, Shoemaker D (2019). Global invasion history of the world's most abundant pest butterfly: a citizen science population genomics study. *Proceedings of the National Academy of Sciences (PNAS) 116*(40), 20015–20024.
Stefanescu, C., Peñuelas, J., & Filella, I. (2003). Effects of climatic change on the phenology of butterflies in the northwest Mediterranean Basin. *Global change biology 9*(10), 1494–1506.
Walker, T. J. (2001). Butterfly migrations in Florida: seasonal patterns and long-term changes. *Environmental entomology 30*(6), 1052–1060.
Williams, C. B. (1930). The migration of butterflies. *The Migration of Butterflies.*

**Plutella xystosella**

Campos, W. G., Schoereder, J. H., & DeSouza, O. F. (2006). Seasonality in neotropical populations of *Plutella xylostella* (Lepidoptera): resource availability and migration. *Population Ecology*, *48*(2), 151158.

Chapman, J. W., Reynolds, D. R., Smith, A. D., Riley, J. R., Pedgley, D. E., & Woiwod, I. P. (2002). High-altitude migration of the diamondback moth *Plutella xylostella* to the UK: a study using radar, aerial netting, and ground trapping. *Ecological Entomology*, *27*(6), 641–650.

Chen, M. Z., Cao, L. J., Li, B. Y., Chen, J. C., Gong, Y. J., Yang, Q., ... & Wei, S. J. (2020). Migration trajectories of the diamondback moth *Plutella xylostella* in China inferred from population genomic variation. *Pest Management Science*.

Furlong, M. J., Wright, D. J., & Dosdall, L. M. (2013). Diamondback moth ecology and management: problems, progress, and prospects. *Annual review of entomology*, *58*, 517–541.

Harcourt, D. G. (1957). Biology of the Diamondback moth, *Plutella maculipennis* (Curt.)(Lepidoptera: Plutellidae), in Eastern Ontario. II. Life-history, behaviour, and host relationships 1. *The Canadian Entomologist*, *89*(12), 554–564.

Shirai, Y. (1995). Longevity, flight ability and reproductive performance of the diamondback moth, *Plutella xylostella* (L.)(Lepidoptera: Yponomeutidae), related to adult body size. *Population Ecology*, *37*(2), 269–277.

Smith, D. B., & Sears, M. K. (1982). Evidence for dispersal of diamondback moth, *Plutella xylostella* (Lepidoptera: *Plutellidae*), into southern Ontario.

Sparks, T. H., Dennis, R. L., Croxton, P. J., & Cade, M. (2007). Increased migration of Lepidoptera linked to climate change. *European Journal of Entomology*, *104*(1), 139.

Williams, C. B. (1930). The migration of butterflies. *The Migration of Butterflies.*

Yang, J., Tian, L., Xu, B., Xie, W., Wang, S., Zhang, Y., ... & Wu, Q. (2015). Insight into the migration routes of Plutella xylostella in China using mt COI and ISSR markers. *PLoS One*, *10*(6), e0130905. Yang, J., Tian, L., Xu, B., Xie, W., Wang, S., Zhang, Y., ... & Wu, Q. (2015). Insight into the migration routes of *Plutella xylostella* in China using mt COI and ISSR markers. PLoS One, 10(6), e0130905.

Yoshimatsu, S. I. (1991). Lepidopterous insects captured on East China Sea from 1981 to 1987. *Japanese Journal of Entomology*, *59*(4), 811–820.

**Vanessa cardui**

Abbot C.H. 1951. A quantitative study of the migration of the painted lady butterfly, Vanessa cardui *L. Ecology* 32**,** 155–171.

Giuliani D, Shields O. 1995. Large-scale migrations of the Painted lady butterfly Vanessa cardui (Lepidoptera: Nymphalidae), in Inyo County, California, during 1991. *Bulletin of the Southern California Academy of Sciences* 94**,** 149–168.

Lokki J, Malmstrom KK, Suomalainen E. 1978. Migration of *Vanessa cardui* new record and *Plutella xylostella* (Lepidoptera) to Spitsbergen in the summer 1978. *Not. Entomol.* 58, 121–123.

Menchetti M, Guéguen M, Talavera G (2019). *Spatiotemporal niche modelling of multigenerational insect migrations. Proceedings of the Royal Society B***.** 286:1910.

Myres, M.T. 1985. A southward return migration of painted lady butterflies, Vanessa cardui, over southern Alberta in the fall of 1983, and biometeorological aspects of their outbreaks into North America and Europe. *Canadian Field Naturalist,* 99**,** 147–155.

Larsen, T. B. (1975). Provisional notes on migrant butterflies in Lebanon. *Atalanta Munnerstadt*, 62, 62–74.

Stefanescu C, Alarcón M, Avila A. 2007. Migration of the painted lady butterfly, Vanessa cardui, to north-eastern Spain is aided by African wind currents. *Journal of Animal Ecology* 76, 888–898.

Stefanescu C, Askew RR, Corbera J, Shaw MR. (2012) Parasitism and migration in southern Palaearctic populations of the painted lady butterfly, *Vanessa cardui* (Lepidoptera: Nymphalidae). *European Journal of Entomology* 109, 85–94.

Stefanescu C, Páramo F, Åkesson S, Alarcón M, Ávila A, Brereton T, Carnicer J, Cassar LF, Fox R, Heliölä J, Hill JK, Hirneisen N, Kjellén N, Kühn E, Kuussaari M, Leskinen M, Liechti F, Musche Regan EC, Reynolds DR, Roy DB, Ryrholm N, Schmaljohann H, Settele J, Thomas CD, van Swaay C, Chapman JW. (2013) Multi-generational longdistance migration of insects: studying the painted lady butterfly in the Western Palaearctic. *Ecography* 36, 474–486.

Stefanescu, C., Soto, D. X., Talavera, G., Vila, R., & Hobson, K. A. (2016). Long-distance autumn migration across the Sahara by painted lady butterflies: exploiting resource pulses in the tropical savannah. *Biology letters*, *12*(10), 20160561.

Suchan T, Talavera G, Sáez L, Ronikier M, Vila R (2018). *Pollen metabarcoding as a tool for tracking long-distance insect migration. Molecular Ecology Resources. 19(1):149-162*

Talavera, G., Bataille, C., Benyamini, D., Gascoigne-Pees, M., & Vila, R. (2018). Round-trip across the Sahara: Afrotropical Painted Lady butterflies recolonize the Mediterranean in early spring. *Biology letters*, *14*(6), 20180274.

Talavera, G., & Vila, R. (2017). Discovery of mass migration and breeding of the painted lady butterfly Vanessa cardui in the Sub-Sahara: the Europe–Africa migration revisited. *Biological Journal of the Linnean Society*, *120*(2), 274-285.

Tilden, J.W. 1962. General characteristics of the movements of *Vanessa cardui* (L.). *Journal of Research on the Lepidoptera* 1, 43–49.

Vandenbosch, R. 2003. Fluctuations of *Vanessa cardui* butterfly abundance with El Niño and Pacific Decadal Oscillation climatic variables. *Global Change Biology,* 9, 785–790.

Williams, C. B. (1930). The migration of butterflies. *The Migration of Butterflies.*

Williams CB. (1970) The migrations of the painted lady butterfly, *Vanessa cardui* (Nymphalidae), with special reference to North America. *Journal of the Lepidopterists' Society* 24, 157–175.

**Zerene cesonia**

Chu, J. P., & Sportiello, M. (2007). *Butterfly research in Boulder County, Colorado 2004-2007* (Doctoral dissertation, Colorado State University. Libraries).

Chu, J. (2009). Inventories of Butterflies in Boulder County. *Unpublished report for Boulder County Parks and Open Space, Boulder, CO.*

Chowdhury, S., Fuller, R. A., Dingle, H., Chapman, J. W., & Zalucki, M. P. (2021). Migration in butterflies: a global overview. *Biological Reviews*, *96*(4), 1462-1483.

Hobson, K. A., Kusack, J. W., & Mora-Alvarez, B. X. (2021). Origins of six species of butterflies migrating through northeastern Mexico: New insights from stable isotope (δ2H) analyses and a call for documenting butterfly migrations. *Diversity*, *13*(3), 102.
