## Supplementary material for "Migratory behavior is positively associated with genetic diversity in butterflies": Figure S2

**Figure S2.** GenomeScope *k-mer* profile plot of the 97 butterfly species dataset showing the fit of the GenomeScope model (black) to the observed *k-mer* frequencies (blue). For each species, the plot shown displayed higher minimum model fit among *k-mer* size 15, 17, 19, 21, 23, 25. The *k-mer* size used and NCBI accession number for the genomic reads used are indicated after species name.

***Agraulis vanillae*** ERR5235460 kmer25

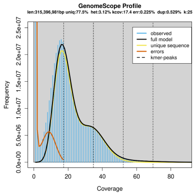


***Agraulis vanillae*** SRR8883910 kmer25

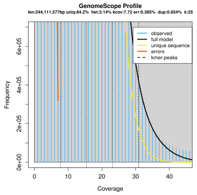


***Agrotis ipsilon*** SRR6432899 kmer25

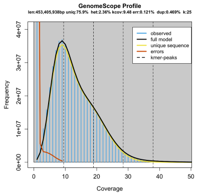


***Agrotis ipsilon*** SRR8103939 kmer21

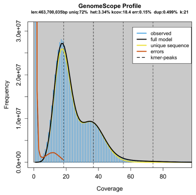


***Amyelois transitella*** SRR1946629 kmer21

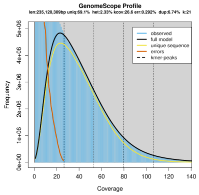


***Amyelois transitella*** SRR9117089 kmer19

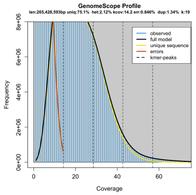


***Amyelois transitella*** SRR9117090 kmer25

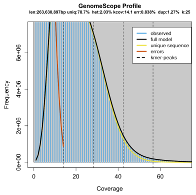


***Aporia crataegi*** SRR7948941 kmer25

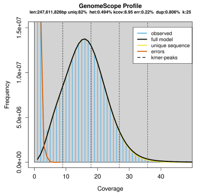


***Aporia largeteaui*** SRR13958260 kmer23

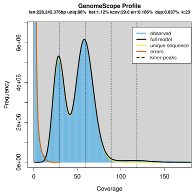


***Arctia plantaginis*** ERR3909543 kmer21

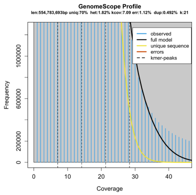


***Arctia plantaginis*** ERR3909544 kmer23

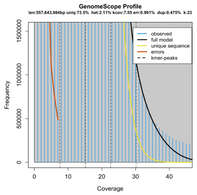


***Baronia brevicornis*** SRR8954515 kmer25

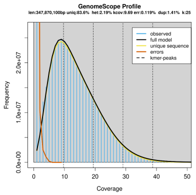

***Bicyclus anynana*** ERR1102678 kmer25

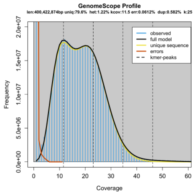


***Bicyclus anynana*** ERR1750945 kmer21

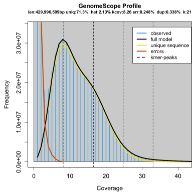


***Bicyclus anynana*** ERR1750946 kmer25

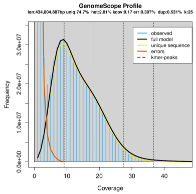


***Bombyx mandarina*** SRR6111375 kmer25

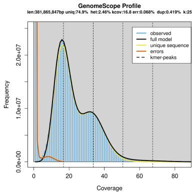


***Bombyx mandarina*** SRR6111379 kmer21

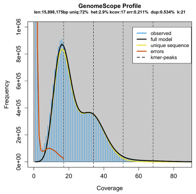


***Bombyx mandarina*** SRR6111382 kmer25

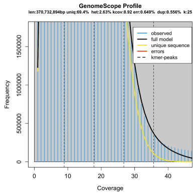


***Bombyx mori*** SRR6111399 kmer25

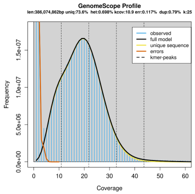


***Bombyx mori*** SRR6111404 kmer25

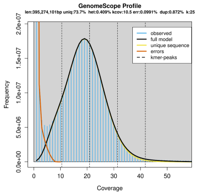


***Calycopis cecrops*** SRR3397505 kmer17

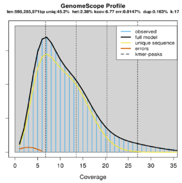


***Charaxes varanes*** SRR5175869 kmer25

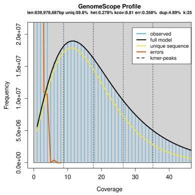


***Charaxes varanes*** SRR8238571 kmer25

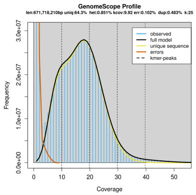


***Chilo suppressalis*** SRR8238596 kmer25

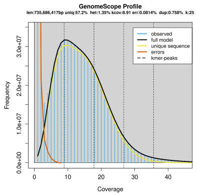

***Chilo suppressalis*** SRR8238690 kmer25

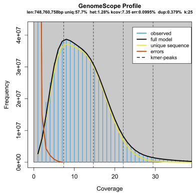


***Colias croceus*** SRR10397971 kmer21

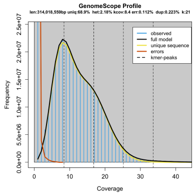


***Colias croceus*** SRR10397986 kmer21

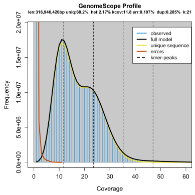


***Colias croceus*** SRR10397988 kmer25

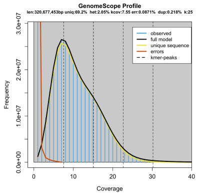


***Colias eurytheme*** SRR11315817 kmer21

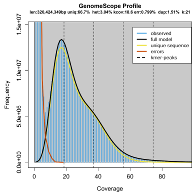


***Colias eurytheme*** SRR14168747 kmer23

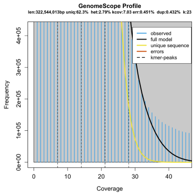


***Colias philodice*** SRR14168755 kmer21

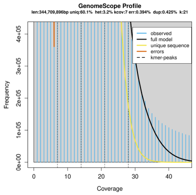


***Colias philodice*** SRR14305386 kmer25

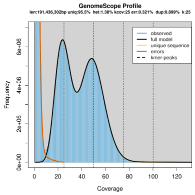


***Cressida cressida*** SRR8954511 kmer17

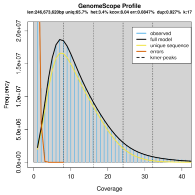


***Cyclargus thomasi*** SRR6727422 kmer25

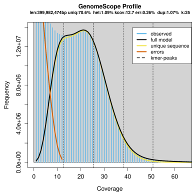


***Danaus chrysippus*** ERR3773468 kmer21

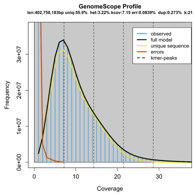


***Danaus chrysippus*** ERR3773475 kmer21

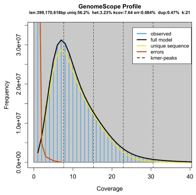

***Danaus chrysippus*** ERR3773477 kmer25

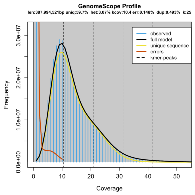


***Danaus eresimus*** SRR1552522 kmer23

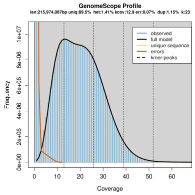


***Danaus eresimus*** SRR1552523 kmer25

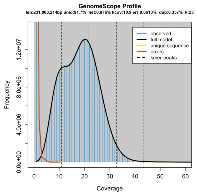


***Danaus erippus*** SRR1552520 kmer25

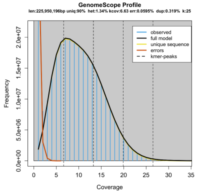


***Danaus erippus*** SRR1552521 kmer25

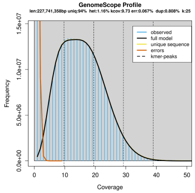


***Danaus gilippus*** SRR1552524 kmer23

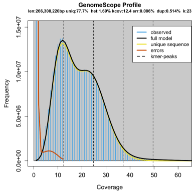


***Danaus gilippus*** SRR1980588 kmer23

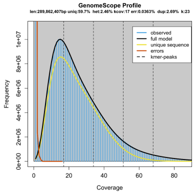


***Danaus melanippus*** SRR3493384 kmer17

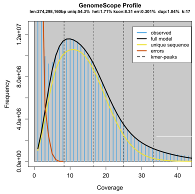


***Danaus plexippus (Australia)*** SRR1552518 kmer21

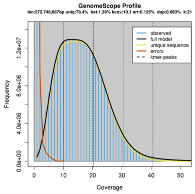


***Danaus plexippus (Florida)*** SRR1552001 kmer21

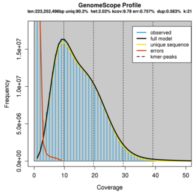


***Danaus plexippus (Hawaii)*** SRR1552232 kmer21

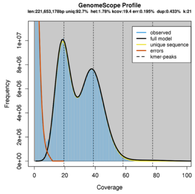


***Danaus plexippus (Massachussets)*** SRR1548578 kmer25

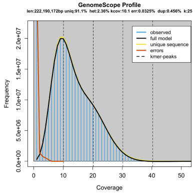

***Danaus plexippus* (New Zealand)** SRR1552236 kmer21

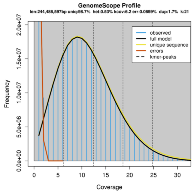


***Danaus plexippus* (Portugal)** SRR1551992 kmer21

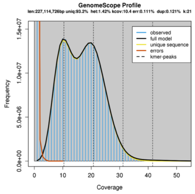


***Danaus plexippus* (Samoa)** SRR1549537 kmer21

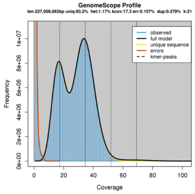


***Depressaria pastinacella*** SRR6984049 kmer25

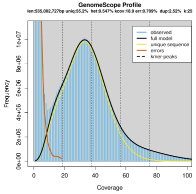


***Depressaria pastinacella*** SRR6984052 kmer25

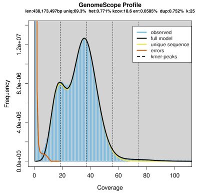


***Depressaria pastinacella*** SRR6984055 kmer23

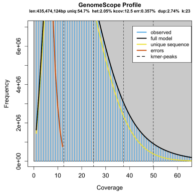


***Doxocopa laurentia*** SRR11638254 kmer25

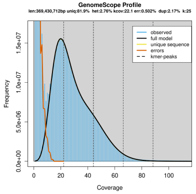


***Dryadula phaetusa*** ERR5235469 kmer25

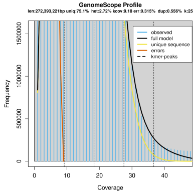


***Ectropis grisescens*** SRR13948480 kmer25

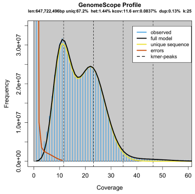


***Ectropis grisescens*** SRR13948481 kmer25

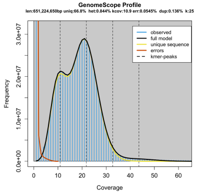


***Ectropis grisescens*** SRR6432897 kmer25

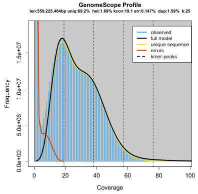


***Elymnias hypermnestra*** SRR12544679 kmer25

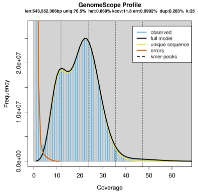

***Elymnias hypermnestra*** SRR12558869 kmer25

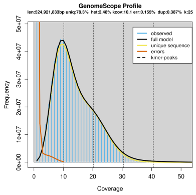


***Eueides isabella*** ERR1051265 kmer25

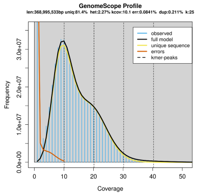


***Eueides isabella*** ERR1051266 kmer25

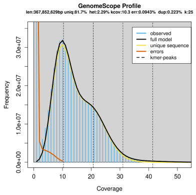


***Eumaeus atala*** SRR13614697 kmer25

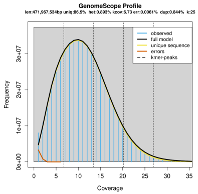


***Eumaeus atala*** SRR13614698 kmer25

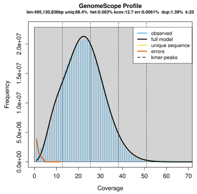


***Eumaeus atala*** SRR6727440 kmer25

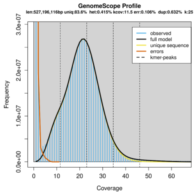


***Hebomoia glaucippe*** SRR13516308 kmer25

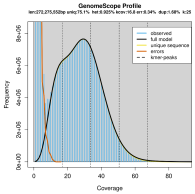


***Heliconius besckei*** SRR3102011 kmer19

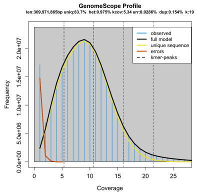


***Heliconius besckei*** SRR3102061 kmer25

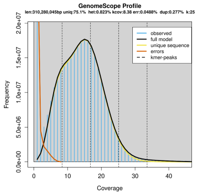


***Heliconius besckei*** SRR3102131 kmer23

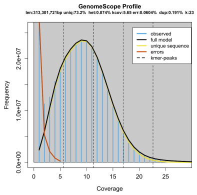


***Heliconius burneyi*** SRR8883892 kmer25

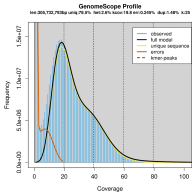


***Heliconius charithonia*** ERR5235470 kmer25

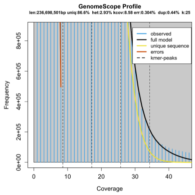

***Heliconius charitonia*** SRR4032025 kmer15

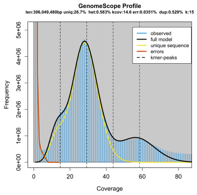


***Heliconius charitonia*** SRR4032026 kmer25

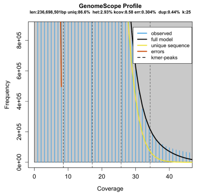


***Heliconius clysonymus*** SRR4032079 kmer17

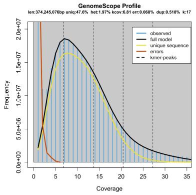


***Heliconius congener*** SRR3102172 kmer19

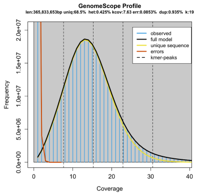


***Heliconius eleuchia*** SRR3102171 kmer25

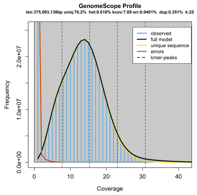


***Heliconius ethilla*** ERR1143576 kmer21

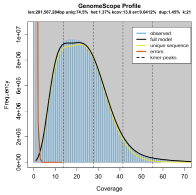


***Heliconius ethilla*** ERR1143577 kmer21

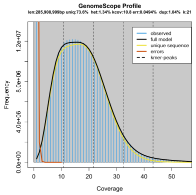


***Heliconius ethilla*** ERR1143578 kmer17

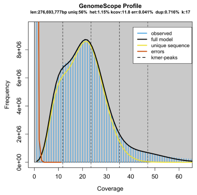


***Heliconius hecalesia*** SRR4031998 kmer19

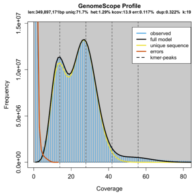


***Heliconius hermathena*** SRR10746200 kmer23

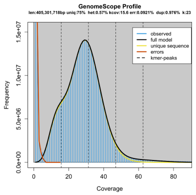


***Heliconius hermathena*** SRR10761151 kmer25

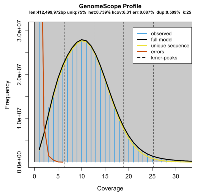


***Heliconius hermathena*** SRR10761171 kmer25

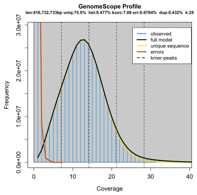

***Heliconius hermathena*** SRR4032035 kmer25

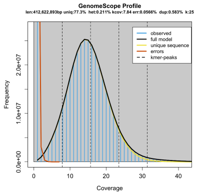


***Heliconius heurippa*** ERR1143584 kmer25

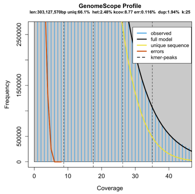


***Heliconius heurippa*** ERR2298267 kmer23

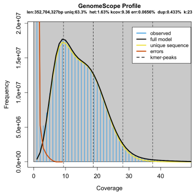


***Heliconius hewitsoni*** SRR3102337 kmer25

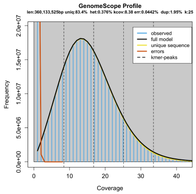


***Heliconius hortense*** SRR4032054 kmer25

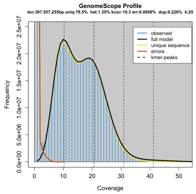


***Heliconius ismenius*** SRR7162647 kmer25

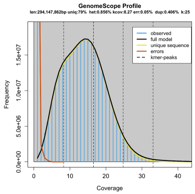


***Heliconius ismenius*** SRR7162649 kmer25

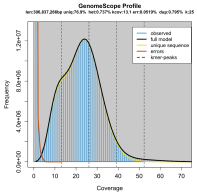


***Heliconius ismenius*** SRR7162650 kmer25

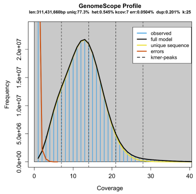


***Heliconius melpomene*** ERR3401274 kmer25

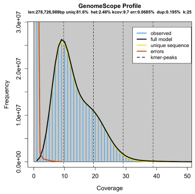


***Hyphantria cunea*** SRR8103944 kmer25

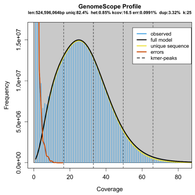


***Hyphantria cunea*** SRR8109446 kmer25

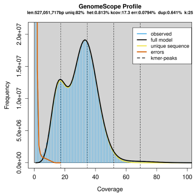


***Hyphantria cunea*** SRR8109447 kmer25

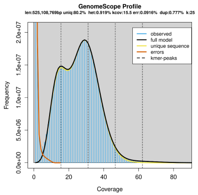

***Jalmenus evagoras*** SRR6679362 kmer25

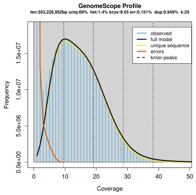


***Junonia oenone*** SRR5182725 kmer25

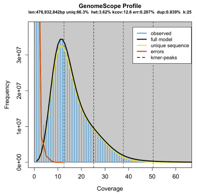


***Kallima inachus*** SRR10276648 kmer21

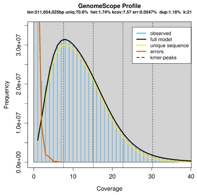


***Lepidochrysops patricia*** SRR6679360 kmer25

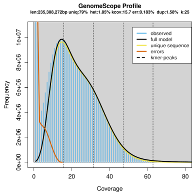


***Leptidea juvernica*** ERR2040009 kmer19

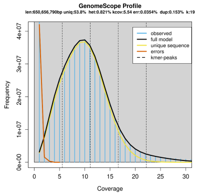


***Leptidea juvernica*** ERR2040010 kmer25

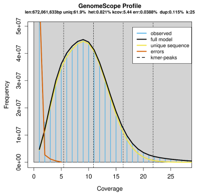


***Leptidea juvernica*** ERR2040015 kmer21

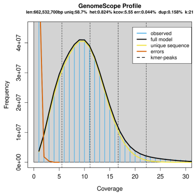


***Leptidea sinapis*** ERR2039865 kmer23

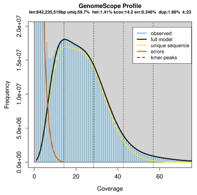


***Leptidea sinapis*** ERR2040021 kmer23

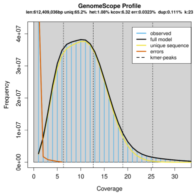


***Leptidea sinapis*** ERR2040025 kmer23

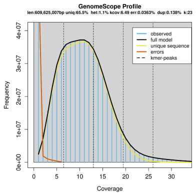


***Lerema accius*** SRR2089774,SRR2089775 kmer17.png

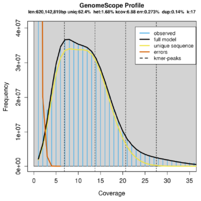


***Limenitis archippus*** SRR9827776 kmer25

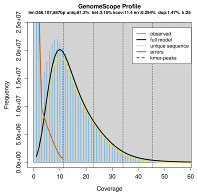

***Limenitis lorquini*** SRR9827807 kmer17

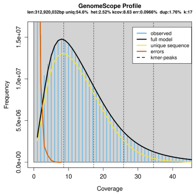


***Limenitis lorquini*** SRR9827808 kmer25

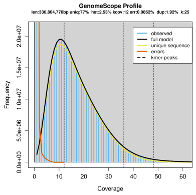


***Lymantria dispar*** SRR13518687 kmer25

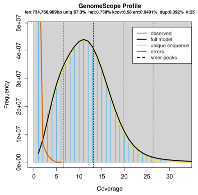


***Lymantria dispar*** SRR13518688 kmer25

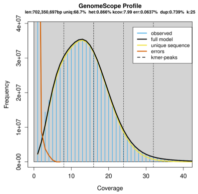


***Maniola jurtina*** SRR9707499 kmer25

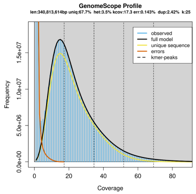


***Ornithoptera priamus*** SRR8954527 kmer25

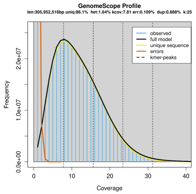


***Ostrinia furnacalis*** SRR6432898 kmer25

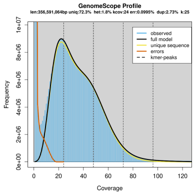


***Ostrinia furnacalis*** SRR8103940 kmer25

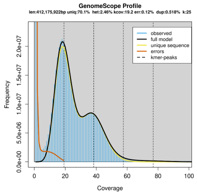


***Ostrinia nubilalis*** SRR12414138 kmer19

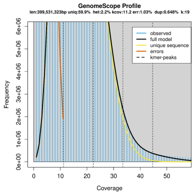


***Ostrinia nubilalis*** SRR12485849 kmer21

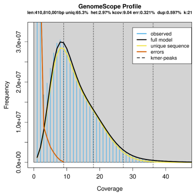


***Ostrinia nubilalis*** SRR8996146 kmer19

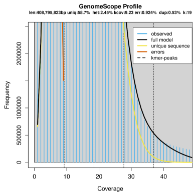


***Ostrinia penitalis*** SRR12485798 kmer19

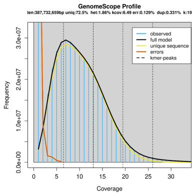

***Ostrinia penitalis*** SRR12485799 kmer25

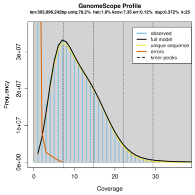


***Ostrinia penitalis*** SRR12485800 kmer25

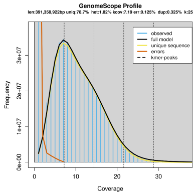


***Ostrinia scapulalis*** ERR2564012 kmer25

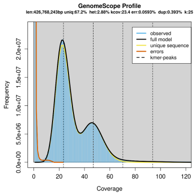


***Papilio aegeus*** SRR10532860 kmer25

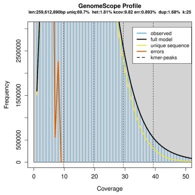


***Papilio aegeus*** SRR10532871 kmer21

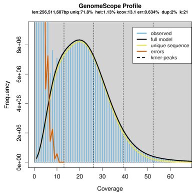


***Papilio aegeus*** SRR10532988 kmer19

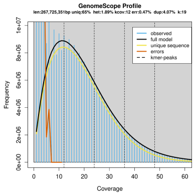


***Papilio aegeus*** SRR10532990 kmer25

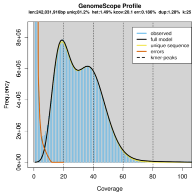


***Papilio alcmenor*** SRR10532966 kmer25

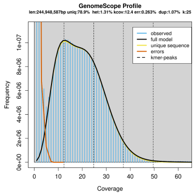


***Papilio ambrax*** SRR10532940 kmer23

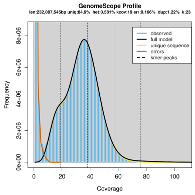


***Papilio ambrax*** SRR10532941 kmer25

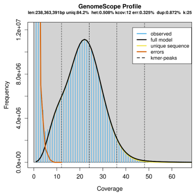


***Papilio ambrax*** SRR10532942 kmer25

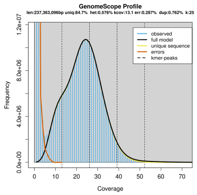


***Papilio ambrax*** SRR10532955 kmer25

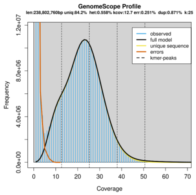

***Papilio aristodemus*** SRR6727429 kmer23

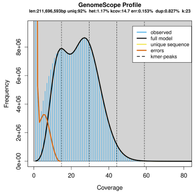


***Papilio ascalaphus*** SRR10532934 kmer25

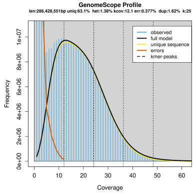


***Papilio ascalaphus*** SRR10532935 kmer25

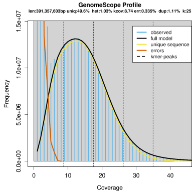


***Papilio ascalaphus*** SRR10532936 kmer25

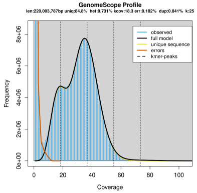


***Papilio bootes*** SRR10532933 kmer23

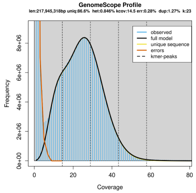


***Papilio castor*** RR10532931 kmer23

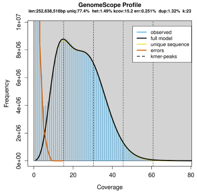


***Papilio castor*** SRR10532932 kmer23

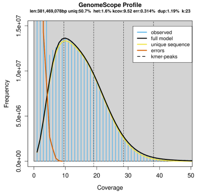


***Papilio gigon*** SRR4342886 kmer23

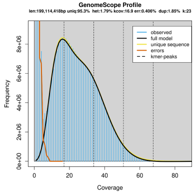


***Papilio glaucus*** SRR1707280 kmer19

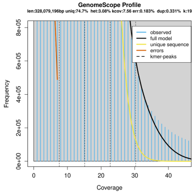


***Papilio iswara*** SRR10532927 kmer25

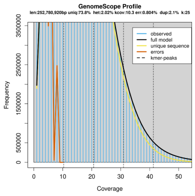


***Papilio joanae*** SRR8954524 kmer19

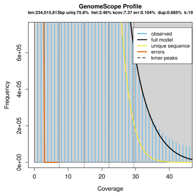


***Papilio machaon*** SRR1777399 kmer17.png

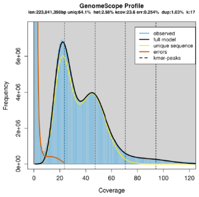

***Papilio macilentus*** SRR10532925 kmer23

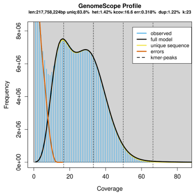


***Papilio memnon*** SRR10532898 kmer21

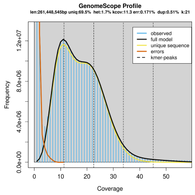


***Papilio memnon*** SRR10532899 kmer25

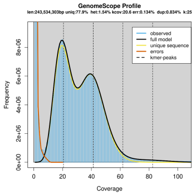


***Papilio memnon*** SRR10532919 kmer25

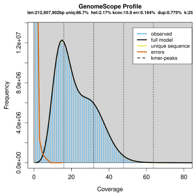


***Papilio memnon*** SRR4341359 kmer25

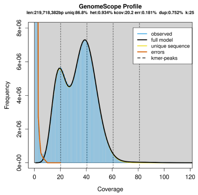


***Papilio nephelus*** SRR10532881 kmer25

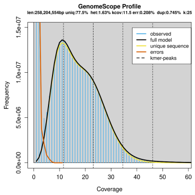


***Papilio nephelus*** SRR10532888 kmer25

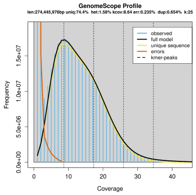


***Papilio nephelus*** SRR10532889 kmer23

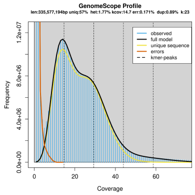


***Papilio nephelus*** SRR10532890 kmer25

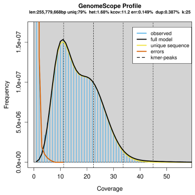


***Papilio oenomaus*** SRR10532879 kmer23

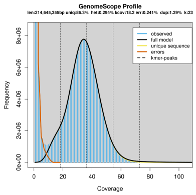


***Papilio phestus*** SRR5879277 kmer25

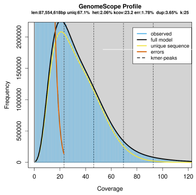


***Papilio phestus*** SRR5879278 kmer23

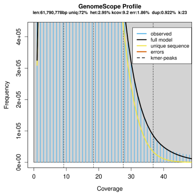

***Papilio phestus*** SRR5879279 kmer23

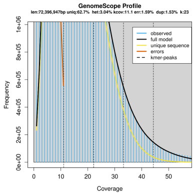


***Papilio polymnestor*** SRR10532876 kmer23

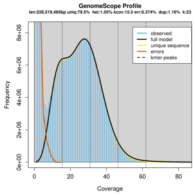


Papilio polymnestor SRR10532877 kmer25

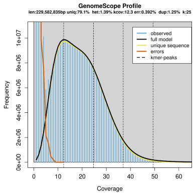


***Papilio polytes*** DRR021666 kmer25

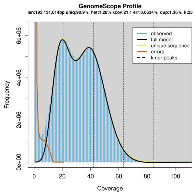


***Papilio polytes*** SRR1107999 kmer25

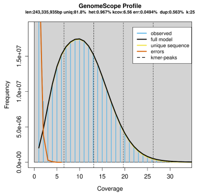


***Papilio polytes*** SRR5879280 kmer25

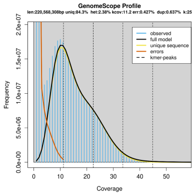


***Papilio polytes*** SRR5879292 kmer25

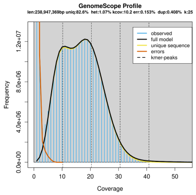


***Papilio polyxenes*** SRR5168526 kmer23

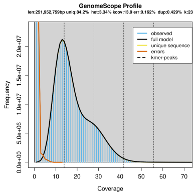


***Papilio protenor*** SRR10532872 kmer25

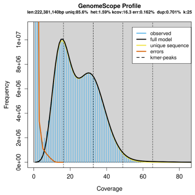


***Papilio protenor*** SRR5879276 kmer23

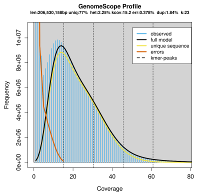


***Papilio protenor*** SRR5879297 kmer19

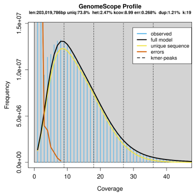


***Papilio sataspes*** SRR10532958 kmer23

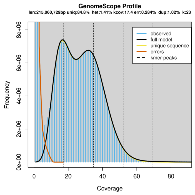

***Papilio sataspes*** SRR10532959 kmer21

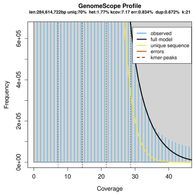


***Papilio sataspes*** SRR10532960 kmer25

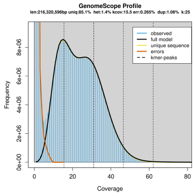


***Papilio thaiwanus*** SRR10532957 kmer23

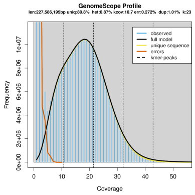


***Papilio tydeus*** SRR10532951 kmer21

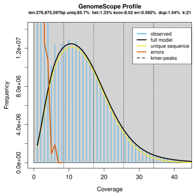


***Papilio tydeus*** SRR10532953 kmer25

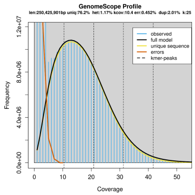


***Papilio tydeus*** SRR10532954 kmer23

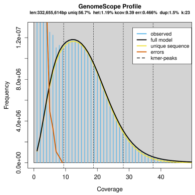


***Papilio tydeus*** SRR10532956 kmer25

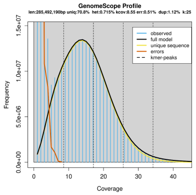


***Papilio xuthus*** SRR11271947 kmer25

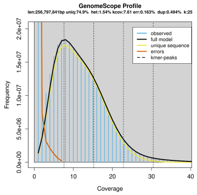


***Papilio xuthus*** SRR1778050 kmer17.png

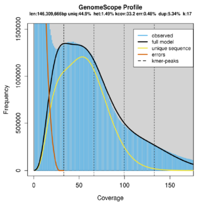


***Papilio xuthus*** SRR2058043 kmer25

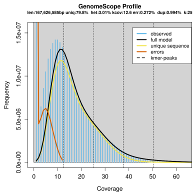


***Papilio zelicaon*** SRR8954529 kmer23

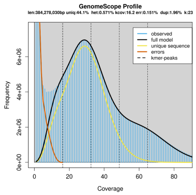


***Pararge aegeria*** SRR1190479 kmer25

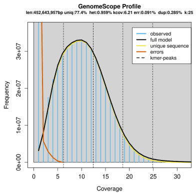

***Pararge aegeria*** SRR7637637 kmer25

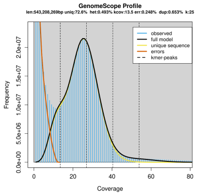


***Pararge aegeria*** SRR7637638 kmer25

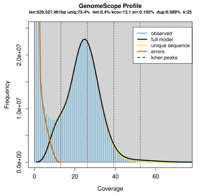


***Parides photinus*** SRR8954530 kmer25

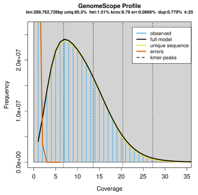


***Pharmacophagus antenor*** SRR8954535 kmer25

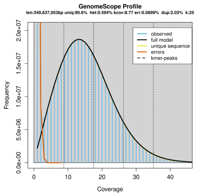


***Phoebis sennae*** SRR3091617 kmer17.png

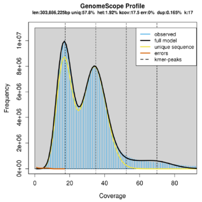


***Pieris macdunnoughi*** ERR5056074 kmer25

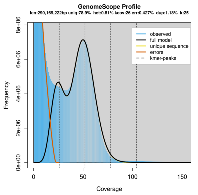


***Pieris napi*** ERR2286688 kmer21

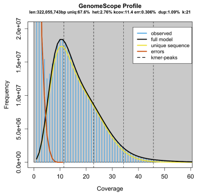


***Pieris napi*** SRR13249445 kmer25

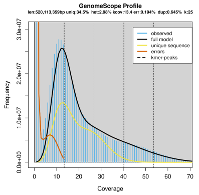


***Pieris rapae*** SRR4339878 kmer25

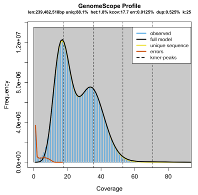


***Plutella xylostella*** SRR6505219 kmer21

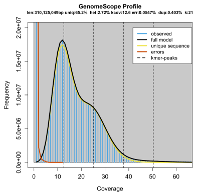


***Polyura narcaeus*** SRR14040768 kmer25

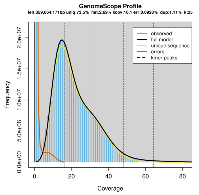


***Proboscis propylea*** SRR12131450 kmer25

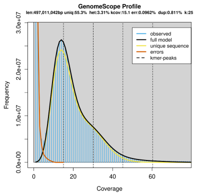

***Vanessa cardui*** A708 kmer21

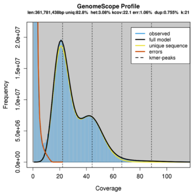


***Vanessa cardui*** D kmer21

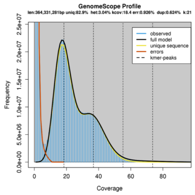


***Vanessa cardui*** E kmer21

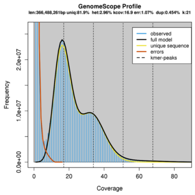


***Vanessa cardui*** VC kmer21

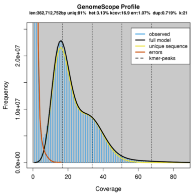


***Vanessa tameamea*** SRR6442215 kmer17.png

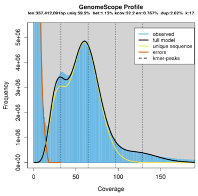


***Zerene cesonia*** SRR11021458 kmer25

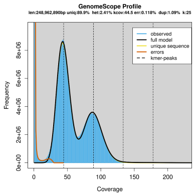


***Zerene cesonia*** SRR11021459 kmer25

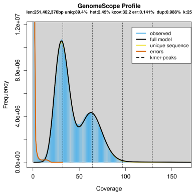


***Zerene eurydice*** SRR11315818 kmer25

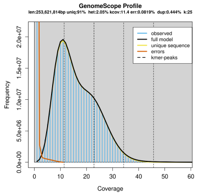


***Zerene eurydice*** SRR14347996 kmer19

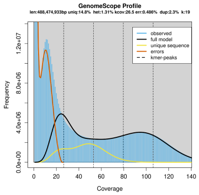


***Zerene eurydice*** SRR14347997 kmer21

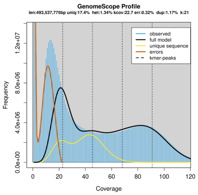
